# Laser Desorption Rapid Evaporative Ionisation Mass Spectrometry (LD-REIMS) demonstrates a direct impact of hypochlorous acid stress on PQS-mediated quorum sensing in *Pseudomonas aeruginosa*

**DOI:** 10.1101/2023.10.11.561961

**Authors:** Rob Bradley, Daniel Simon, Livia Spiga, Yuchen Xiang, Zoltan Takats, Huw Williams

**Affiliations:** Department of Life Sciences, Faculty of Natural Sciences, Imperial College London, London, UK; Department of Metabolism, Digestion and Reproduction, Faculty of Medicine, Imperial College London, London, UK

## Abstract

To establish infections in human hosts, *Pseudomonas aeruginosa* must overcome innate immune generated oxidative stress, such as the hypochlorous acid (HOCl) produced by neutrophils. We set out to find specific biomarkers of oxidative stress through the development of a protocol for the metabolic profiling of *P. aeruginosa* cultures grown in the presence of different oxidants using a novel ionisation technique for mass spectrometry, laser desorption rapid evaporative ionisation mass spectrometry (LD-REIMS). We demonstrated the ability of LD-REIMS to classify samples as untreated or treated with a specific oxidant with 100 % accuracy and identified a panel of 54 metabolites with significantly altered concentrations after exposure to one or more of the oxidants. Key metabolic changes were conserved in *P. aeruginosa* clinical strains isolated from patients with cystic fibrosis lung infections. These data demonstrated that HOCl stress impacted the Pseudomonas Quinolone Signal (PQS) quorum sensing system. Ten 2-alkyl-4-quinolones (AHQs) associated with the PQS system were significantly lower in concentration in HOCl-stressed *P. aeruginosa* cultures, including 2-heptyl-3-hydroxy-4(1*H*)-quinolone (PQS), the most active signal molecule of the PQS system. The PQS system regulates the production of virulence factors, including pyocyanin and elastase, and their levels were markedly affected by HOCl stress. No pyocyanin was detectable and elastase concentrations were significantly reduced in cultures grown with sub-lethal concentrations of HOCl, suggesting that this neutrophil-derived oxidant may disrupt the ability of *P. aeruginosa* to establish infections through interference with production of PQS-associated virulence factors.

**IMPORTANCE:** This work demonstrates that a high-throughput ambient ionisation mass spectrometry method can be used successfully to study a bacterial stress response. Its application to the opportunistic pathogen *Pseudomonas aeruginosa* led to the identification of specific oxidative stress biomarkers, and demonstrated that hypochlorous acid, an oxidant specifically produced by human neutrophils during infection, affects quorum sensing and markedly reduces production of the virulence factors pyocyanin and elastase in this bacterium. This approach has the potential to be widely applicable to the characterisation of the stress responses of bacteria.

## INTRODUCTION

Bacteria encounter oxidative stress in aerobic environments. The innate immune systems of both animals and plants use reactive oxygen species to attack and to attempt to kill invading microbes. Upon phagocytosis of bacteria, the phagocyte or NADPH oxidase of human neutrophils is induced, assembled, and catalyses the formation of the superoxide radical (O ^.-^). This is known as the respiratory burst and is associated with a 10-20-fold increase in O2 consumption (Halliwell & Gutteridge, 2015). The importance of superoxide generated by NADPH oxidase is demonstrated through the susceptibility of people with defective NADPH oxidase to a broad range of microbes, as seen in conditions such as chronic granulomatous disease (CDG) (Marciano et al., 2015). However, superoxide has low reactivity and is membrane impermeable, and so the antimicrobial action of the respiratory burst is thought to be mediated through the conversion of superoxide to stronger oxidants, including H2O2, hypochlorous acid (HOCl) and hypothiocyanous acid (HOSCN) (Lam et al., 2020). O ^.-^ dismutates either spontaneously or catalysed by superoxide dismutase to form H O. HOCl and HOSCN can then be formed by the reaction of H2O2 with chloride or thiocyanate ions respectively, catalysed by myeloperoxidase (MPO). While there exists a large body of evidence to indicate that H2O2 is important for the killing of pathogens by neutrophils (Brudzynski et al., 2011; Palma et al., 2004; Rada & Leto, 2008), the antimicrobial actions of HOCl and HOSCN are relatively poorly studied, despite MPO comprising around 5% of neutrophil proteins (Halliwell & Gutteridge, 2015) and that neutrophil-mediated killing of bacteria has been shown to be MPO-dependent (Klebanoff et al., 2013).

To further understanding of their bactericidal action and the corresponding bacterial protective response, we previously performed a transcriptomic analysis of *P. aeruginosa* grown in HOCl and HOSCN (Farrant et al., 2021). We identified a diverse range of genes required for protection against HOCl and HOSCN, including the transcriptional regulator RclR, a homologue of the *E. coli* HOCl-specific sensor RclR. We demonstrated that RclR regulates the expression of *rclX*, a putative peroxiredoxin, in *P. aeruginosa* and that both RclR and RclX are required specifically for protection against HOCl and HOSCN, with RclX shown to be a HOCl reductase (Nontaleerak et al., 2023). Another key study showed that hypohalous acids, such as HOCl, caused protein aggregation and that *P. aeruginosa* responded by increasing polyphosphate levels, which improved survival by protecting against protein aggregation (Groitl et al., 2017).

We wanted to complement these studies by investigating the metabolic changes that occur following oxidative stress to provide additional mechanistic insights and to identify biomarkers of the stress to provide a way of examining bacterial stress exposure *in vivo* during infection. In parallel we were interested in establishing whether Laser Desorption Rapid Evaporative Ionisation Mass Spectrometry (LD-REIMS) can be used to study a bacterial stress response.

REIMS is a novel ambient ionisation mass spectrometry technique in which rapid heating of a sample generates a sample aerosol containing gas phase ions of metabolites and biological molecules that can be analysed to provide real time, semi-quantitative metabolic profiling. Originally developed as an intra-operative, in situ method for tumour margin detection, REIMS has the major advantage of requiring little-to-no sample preparation (Schäfer et al., 2009). Rapid heating can be achieved through the application of electrical diathermy, for example in the form of a heated electrosurgical knife, or by use of a laser, as in LD-REIMS. Previously, the Takats group used electrical diathermy REIMS to characterise the metabolomes of *P. aeruginosa* clinical isolates by analysing colonies grown on agar plates, successfully classifying isolates to a strain level with 81 % accuracy and identifying a number of *P. aeruginosa* metabolites (Bardin et al., 2018). LD-REIMS has been used successfully for metabolic fingerprinting of biofluids and for screening of libraries for synthetic biology applications, able to detect a panel of diverse biological molecules in a yeast model, demonstrating its potential for broad applicability to metabolite detection (Cameron et al., 2021; Gowers et al., 2019; Plekhova et al., 2021; Van Meulebroek et al., 2020; Wijnant et al., 2020).

The aim of this study was to determine whether LD-REIMS can be used as a high throughput approach to study bacterial stress responses. We wanted first to test whether LD-REIMS can distinguish untreated from oxidatively stressed *P. aeruginosa*, and subsequently whether it can be used to reliably identify changes in the oxidant specific biomarkers in this bacterium, as well as providing mechanistic insights into the response to oxidative stress.

## METHODS

### Bacterial strains & growth conditions

The *P. aeruginosa* lab strain PA14 was used for all experiments unless otherwise stated. The transposon mutant *pqsE* came from a *P. aeruginosa* transposon mutant library (Miyata et al., 2006). The clinical isolates C1, C2, C3, C9, C13, C14, C25, C28 and C50 were supplied by the Royal Brompton Hospital (Table S1). All liquid cultures were grown in LB medium (5 g/L NaCl, 5 g/L yeast extract, and 10 g/L tryptone) at 37 °C, shaking at 700 rpm.

### Preparation of oxidants for bacterial stress

**(i) sodium hypochlorite (HOCl)** HOCl solutions were prepared by diluting NaOCl (Sigma) directly into the media. The molar concentration of OCl^-^ was determined using iodometric titration. Potassium iodide (KI) and hydrochloric acid (HCl) were added to NaOCl, and the resulting solution titrated to a colourless end-point using sodium thiosulphate (Na2S2O3) and a starch solution indicator. **(ii) hypothiocyanous acid (HOSCN):** HOSCN solutions were generated as described in (Chandler et al., 2013) with modifications. Briefly, a solution containing 6.5 mM potassium thiocyanate (KSCN, Fluka) and 6 U/mL lactoperoxidase (LPO, Sigma) was prepared in LB. Aliquots of 1 mM H2O2 were added over 5 minutes, followed by the addition of 100 U/mL catalase (Sigma). To remove proteins, the solution was centrifuged at 4000 rpm for 10 minutes using a 10 kDa filter (Ambion). The concentration of HOSCN was determined by monitoring the loss of signal at 412 nm (ε412 nm = 14,150 M-1 cm-1) that occurs when HOSCN reacts with 5-thio-2-nitrobenzoic acid (TNB) (Nagy et al., 2009). **(iii) hydrogen peroxide (H2O2) solutions:** Solutions were prepared by diluting 30% w/w in H2O (9.79 M, Sigma) directly into the media. **(iv) methylglyoxal (MGO) solutions:** MGO solutions were prepared by diluting MGO (Sigma, 40% in H2O) directly into the media of choice. **(v) methyl viologen (MV) solutions:** MV solutions were prepared by diluting MV (Sigma, 10 mM) directly into the media of choice.

### Preparation of liquid-culture grown bacterial samples for LD-REIMS analysis

LB medium was inoculated with *P. aeruginosa* in a 96 deep-well plate and cultured overnight at 37 °C, shaking at 700 rpm on a THERMOstar microplate incubator and shaker. Each replicate was an independent culture. The overnight cultures were subcultured 1:50 into fresh media spiked with the oxidant of interest (190 µL) in a 96 well-plate and incubated at 37 °C, shaking at 700 rpm. Once mid-log growth phase was achieved, the cultures were transferred to a V-shaped-bottom 96 well-plate (Sigma) and centrifuged at 3270 x *g* for 20 minutes at 4 °C. The supernatant was removed, the pellets resuspended in sterile dH2O (100 µL) and again centrifuged at 3723 x g for 20 minutes, at 4 °C. The supernatant was removed, and the plate containing the bacterial cell pellet was introduced into the LD-REIMS plate reader for analysis.

### LD-REIMS analysis

LD-REIMS analysis was performed by adapting the methodology described by Simon et al., 2023. The pellets were subject to laser ablation using an Opolette 2940 laser system (Opotek). One laser burn was performed per pellet, with a fluence of 5 J/cm^2^ and a laser firing duration of 3 s. Samples were analysed at 3 s intervals. The aerosol produced was transported by vacuum through polytetrafluorethylene tubing (3.2 mm O.D., 1.6 mm I.D.) connected to the REIMS interface through a T-shaped piece, also permitting inflow of isopropanol (IPA) at a flow rate of 0.1 mL/min, as described by Jones et al., 2019. All mass analysis was carried out in negative ionisation mode using a Xevo G2-S QToF instrument with the cone voltage 40 V, heater bias voltage 80 V and *m/z* scan range from 50 to 1200 Da. The REIMS instrument was calibrated in sensitivity mode according to the manufacturer’s standard instructions (Waters Corporation), using 0.1 mM sodium formate solution in isopropanol and water (90/10, v/v) at a flow rate of 0.1 mL/min.

### Data processing and statistical analysis

Peak picking was performed using an in-house workflow based in R and Python and developed by members of the Takats group. In summary, first the raw data were background subtracted and lock mass corrected using one of three known *P. aeruginosa* metabolites as standards depending on the mass range of focus (m/z = 199.1698 / 529.3382 / 716.5231). Fast Fourier Transform filtering was applied to remove high frequency noise in the Fourier domain and the data peak-picked and mass recalibrated using the R package MALDIquant (Gibb & Strimmer, 2012). The resulting data was saved in .csv format, ready for post-processing. Multivariate statistical analysis was performed on the data using the MetaboAnalyst 4.0 platform (Chong et al., 2018). Sum normalisation, logarithmic transformation and Pareto scaling were applied. Principal component analysis (PCA) and orthogonal projections to latent structures discriminant analysis (OPLS-DA) were used for data visualization and support vector machines (SVMs) for classification and feature selection. SVM models were validated using F1 to measure model accuracy, calculated from the model precision and recall.

### LC-MS/MS analysis

*P. aeruginosa* samples were prepared for LC-MS analysis by subculturing overnight cultures 1:50 into fresh LB medium, untreated or spiked with HOCl (3.2 mM). Once an OD600 of 1.0 was achieved, 100 µL of each culture was transferred to a microcentrifuge. 100 µL of Hank’s Balanced Salt Solution (HBSS, Gibco), was added and the mixture incubated at room temperature for 30 minutes. The mixture was then centrifuged at 3270 x *g* for 5 minutes at 4 °C, the supernatant removed, the remaining pellet washed using 1 mL ice-cold ammonium acetate solution (150 mM) in LC-MS grade H2O, centrifuged at 3270 x *g* for 5 minutes at 4 °C. The wash was repeated four more times. After the final supernatant removal, the pellet was stored at -80 °C. LC-MS/MS analysis was performed by the National Phenome Centre (NPC) service using the lipidomics protocol in negative ion mode (see Lewis et al., 2022). Prior to analysis, the pellets were thawed and resuspended in ammonium acetate solution (150 mM). The final ratio of sample to IPA prior to injection was 1:4, as described in the protocol for human serum samples.

### Virulence factor assays

*P. aeruginosa* strains were grown overnight in 5 ml of LB broth in a 50 mL centrifuge tube (Falcon) and incubated overnight at 37°C at 180 rpm, and pyocyanin and elastase measured in the *P. aeruginosa* culture supernatants as previously described (Essar et al., 1990; Filloux & Ramos, 2014).

### Data availability

The data described in this article are openly available on the MassIVE data repository at ftp://MSV000093056@massive.ucsd.edu.

## RESULTS

### 1. Development of a LD-REIMS protocol to study the response of *P. aeruginosa* to oxidative stress

As REIMS has previously only been used to analyse bacterial colonies grown on agar, our first aim was to develop a protocol to analyse stressed, exponentially growing liquid cultures, with minimal disruption of cellular physiology between sampling and analysis, to investigate the metabolic changes occurring during the adaptive response of *P. aeruginosa* to oxidative stress. We grew *P. aeruginosa* cultures in a 96-well microtiter plate format with or without the oxidative stress to an exponential phase of OD600 ∼0.8. The cells were then pelleted and washed with 150 mM ammonium acetate before transfer to the high throughput LD-REIMS autosampler (Fig. 1).

**Figure 1:**
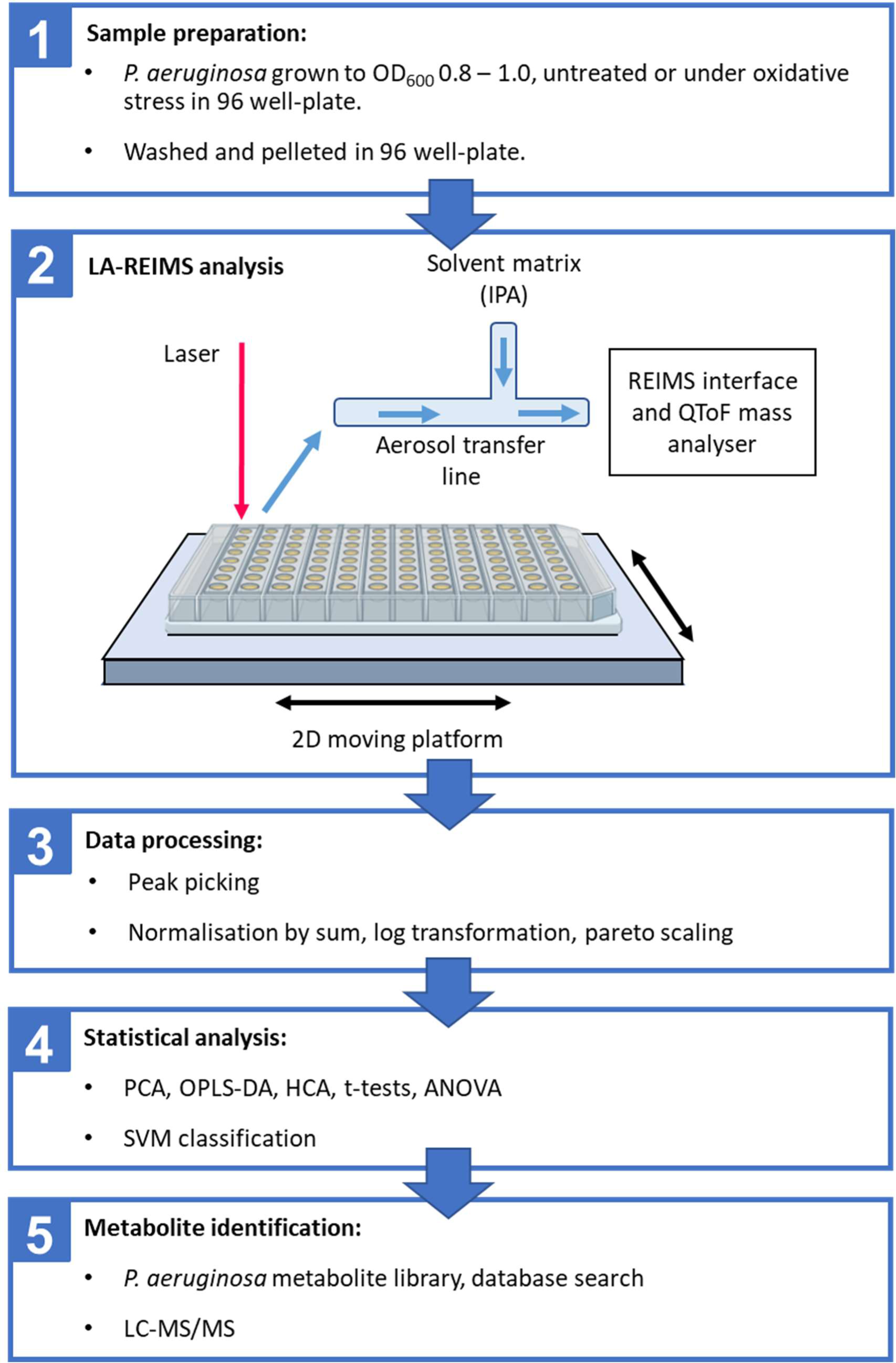
Protocol for high throughput LD-REIMS analysis of P. aeruginosa exposed to oxidative stresses. 1) P. aeruginosa samples were grown in a 96-well plate, untreated or treated following balanced growth in exponential phase. This was achieved by inoculating 10 μL of P. aeruginosa stationary phase cultures into 200 μL of LB to give an initial OD600 of approximately 0.05 followed by growth for 4 hours to an OD600 of 0.8. The cultures were then pelleted, the pellets washed with ammonium acetate and pelleted again. The time between starting to harvest the cultures and LD-REIMS analysis of the pellets was standardised to 60 +/-10 min. 2) Rapid heating and evaporation of the surface of each pellet was performed by moving the 96 well-plate under a CO2 laser using a 2D moving platform. The resulting aerosol was transported to the LD-REIMS interface and analysed using a QToF mass analyser. 3) Once the analysis was complete, the data was first processed using a peak picking script. 4) Statistical analysis was performed on the processed data to determine which spectral features were significantly different between treated and untreated samples. Support vector machine (SVM) classification models were used to classify samples depending on treatment. 5) Metabolite identification was performed using a previous P. aeruginosa metabolite database generated in our lab and online databases. A selection of the most significantly impacted metabolites had their identity confirmed using LCMS/MS.

A typical LD-REIMS spectrum of *P. aeruginosa* samples prepared using this protocol (Fig. 2) showed a similar range of features to those described previously for analysis of *P. aeruginosa* colonies (Bardin et al., 2018), with predominantly 2-alkyl-4-quinolones (AHQs) associated with the PQS quorum sensing (QS) system detected in the range from *m/z* 240 – 320, rhamnolipids from *m/z* 330 – 710 and phospholipids from 600 – 900 *m/z*.

**Figure 2:**
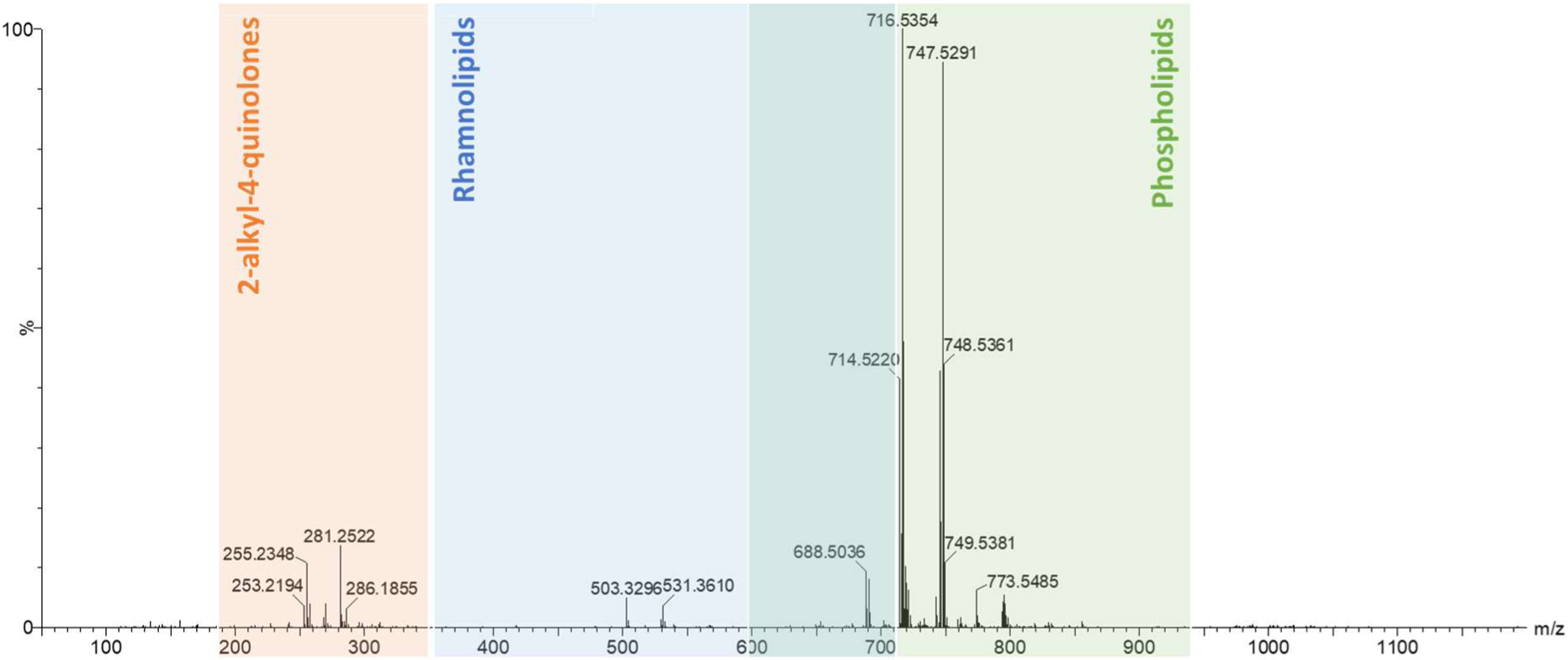
Typical mass spectrum of untreated P. aeruginosa obtained using the LD-REIMS protocol outlined in Fig. 1. The three main biomolecular regions are highlighted: 2-alkyl-4 -quinolones (AHQs) at m/z 240-320, rhamnolipids at m/z 330-710 and phospholipids at m/z 600-900. The main classes of molecule previously identified in LD-REIMS analysis of P. aeruginosa colonies are found here.

### 2. LD-REIMS analysis can distinguish between untreated and oxidatively stressed *P. aeruginosa* cultures

We then tested our LD-REIMS protocol for the detection of metabolic differences between untreated *P. aeruginosa* and cultures stressed by growth in the presence of the oxidants hypochlorous acid (HOCl), hypothiocyanous acid (HOSCN) and hydrogen peroxide (H2O2), the superoxide generator methyl viologen (MV) and the reactive electrophilic compound methylglyoxal (MGO) that has been shown to promote oxidative stress (Figarola et al., 2014). One non-lethal concentration was chosen for each oxidant which led to a one-hour growth lag of the treated cultures compared to untreated. This criterion was chosen to provide evidence that the bacteria were responding to the stress, adapting, and then resuming growth in its presence (Fig. S1). Ten biological replicates were analysed for each of the 6 sample classes (untreated, HOCl, HOSCN, H2O2, MGO and MV).

Given the complexity of the LD-REIMS spectra across the different conditions we first used principal component analysis (PCA), a dimensionality reduction technique that condensed the data and allowed us to visualize the high-level variance patterns in score plots (Fig. 3). Clear clustering of the untreated and treated classes can be observed for each oxidant, indicating that growth in their presence causes substantial metabolic divergence in the bacteria for the biomolecules detected by LD-REIMS.

**Figure 3:**
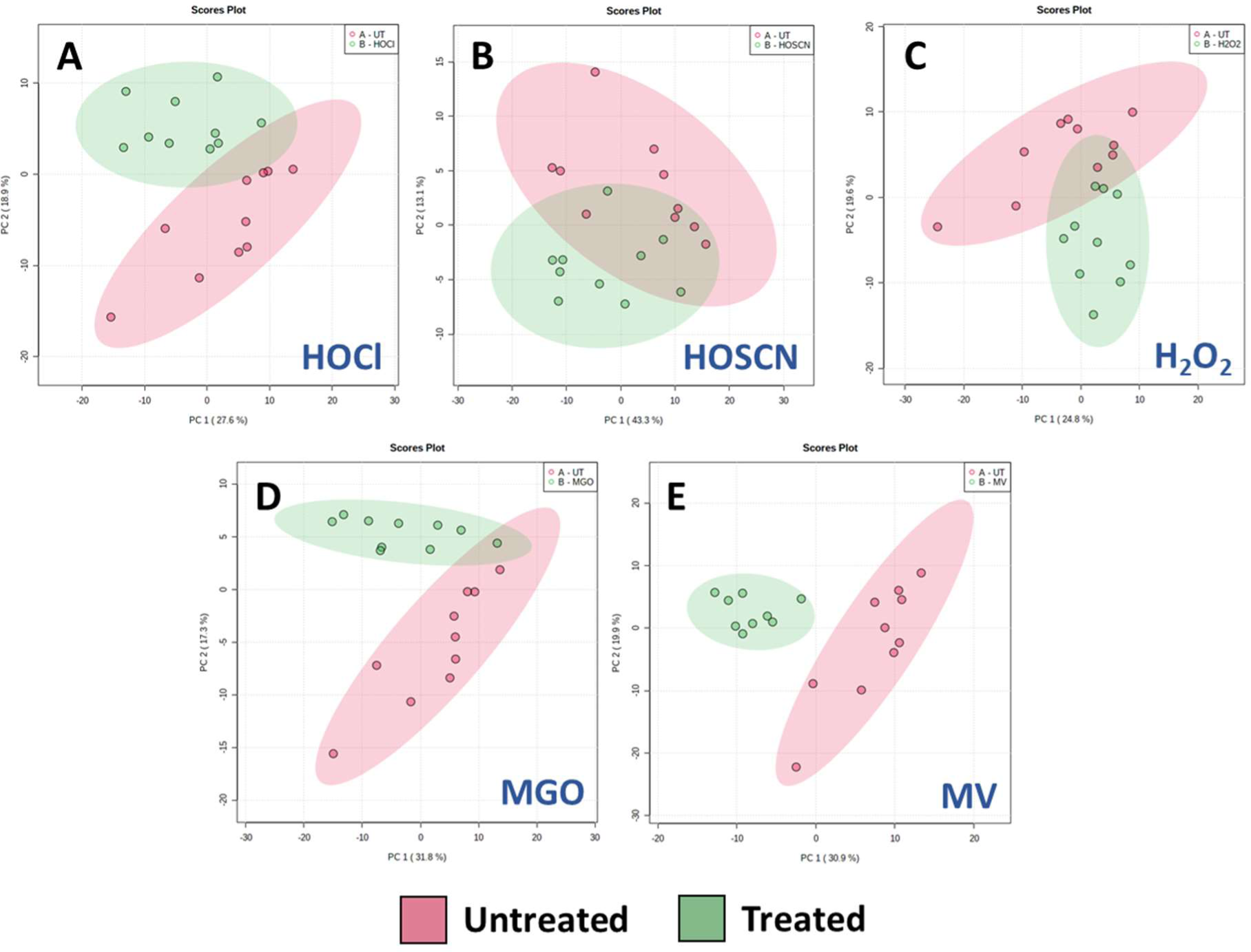
Statistical analysis of P. aeruginosa mass spectra data obtained by LD-REIMS by a two-group PCA of untreated and treated P. aeruginosa samples (n=10); (a) 3.2 mM HOCl, (b) 0.35 mM HOSCN, (c) 60 mM H2O2; (d) 1.8 mM MGO (e) 0.6 mM MV. Statistical analyses were completed on the 50 to 1200 m/z range, after background subtraction and mass drift correction, using the MetaboAnalyst 5.0 platform. The plots all represent principal components one and two. Shaded areas show 95% confidence intervals of the sample groups. Separation between untreated and treatment samples is clear for all oxidants.

We then employed orthogonal projection to latent structures discriminant analysis (OPLS-DA) to confirm the discriminative power of LD-REIMS seen with PCA. OPLS-DA is a supervised modelling method, meaning the class labels are provided and the model identifies the variations in the dataset that are most able to predict the given classes. Whilst this approach provides greater power for class differentiation than PCA, it can be prone to overfitting, meaning the model fits the training data too well and cannot be generalised to new datasets. As one of the aims of this study is to identify biomarkers of oxidative stress that are universal to *P. aeruginosa* samples, it is particularly important to avoid overfitting. Therefore, we cross-validated the OPLS-DA models using the Q^2^ coefficient, a statistic that describes the predictive quality of the model, providing a measure of its reliability. Our analysis generated a Q^2^ value greater than 0.5 for each oxidant (Fig. 4), a commonly employed threshold for validation that suggests the model is less likely to be overfitted (Triba et al., 2015).

**Figure 4:**
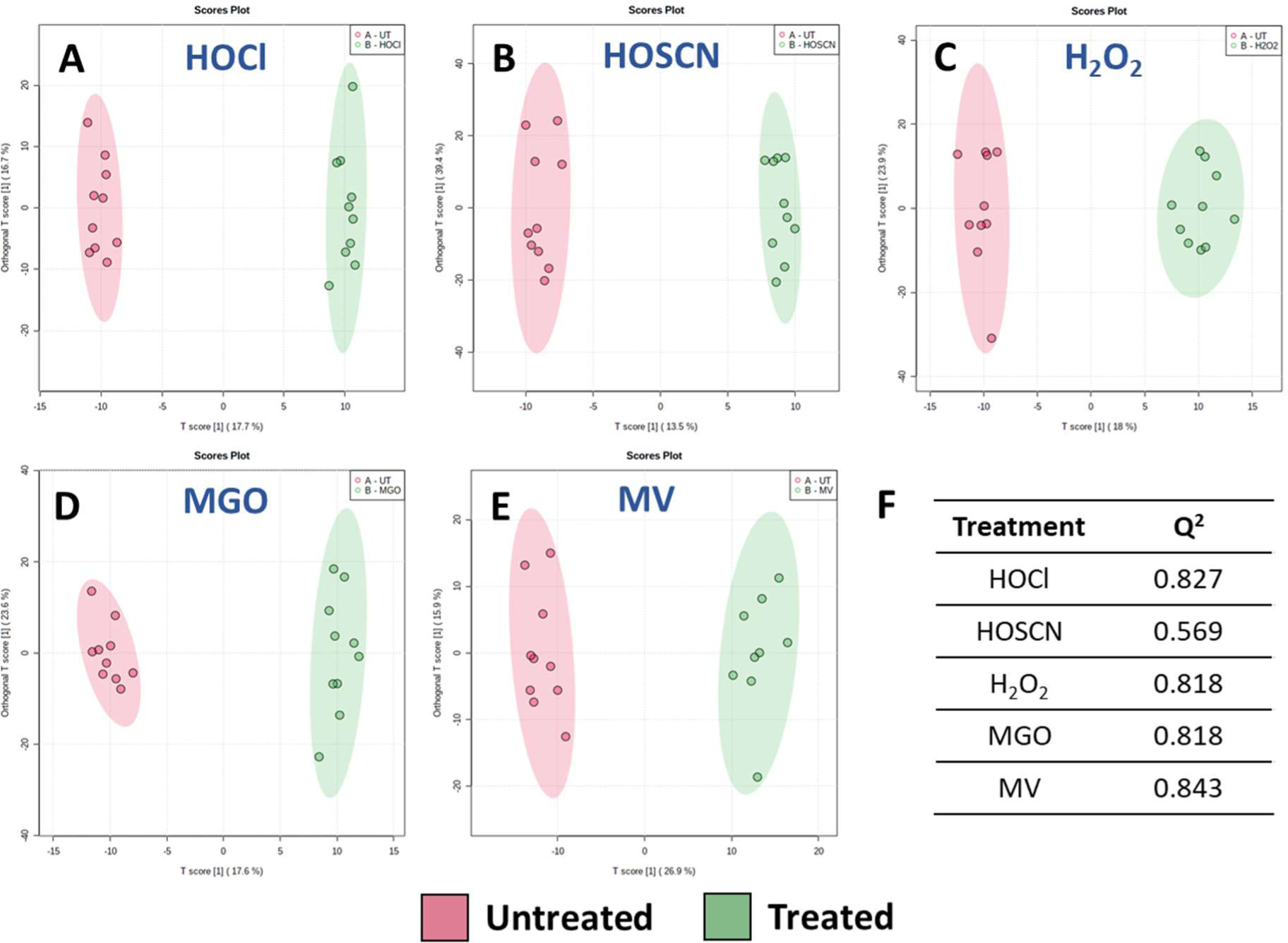
Statistical analysis of P. aeruginosa mass spectra data obtained by LD-REIMS by OPLS-DA analysis. Score plots generated based on OPLS-DA of untreated and treated P. aeruginosa samples (n=10); (a) 3.2 mM HOCl, (b) 0.35 mM HOSCN, (c) 60 mM H2O2; (d) 1.8 mM MGO (e) 0.6 mM MV. Statistical analyses were completed on the 50 to 1200 m/z range, after background subtraction and mass drift correction, using the MetaboAnalyst 5.0 platform. (f) The model is able to separate the untreated and treated samples for each oxidant, cross-validated by the high Q2 coefficient for each plot (Triba et al., 2014).

To provide further confirmation that the model was not overfitted, we tested the reproducibility of the data by performing an identical experiment on new cultures with the same level of biological replication (ten independent cultures) and then comparing the two independent datasets. OPLS-DA generated Q^2^ coefficients similar to the single-day analysis, indicating the discriminative power of the analysis was retained over the two days, despite a reduction in separation in PCA scores plots for HOSCN, H2O2 and MGO that can be attributed to a batch effect (Figs S2 and S3). Overall, these analyses show that LD-REIMS can discriminate clearly and consistently between untreated and oxidatively stressed bacteria for a range of oxidants.

### 3. Can LD-REIMS reliably classify *P. aeruginosa* based on stress exposure?

We next wanted to determine whether LD-REIMS can reliably classify samples based on the specific stress they have been exposed to. To achieve this, we first split the merged dataset at random into a training set and a test set, in an 80:20 ratio. We then used the training set to build a support vector machine (SVM) for classification and validated this model using k-fold cross-validation (k=10). SVMs are commonly employed for large metabolomic datasets due to their resistance to outliers and ability to handle noisy data (Mahadevan et al., 2008). Using the model, the samples were classified depending on oxidant exposure with 100 % accuracy (Fig 5). The model was then tested using the test data set and the samples were again classified with 100 % accuracy, demonstrating that LD-REIMS analysis provides sufficient discriminatory metabolic information to classify samples as treated with a specific oxidant with very high confidence.

**Figure 5:**
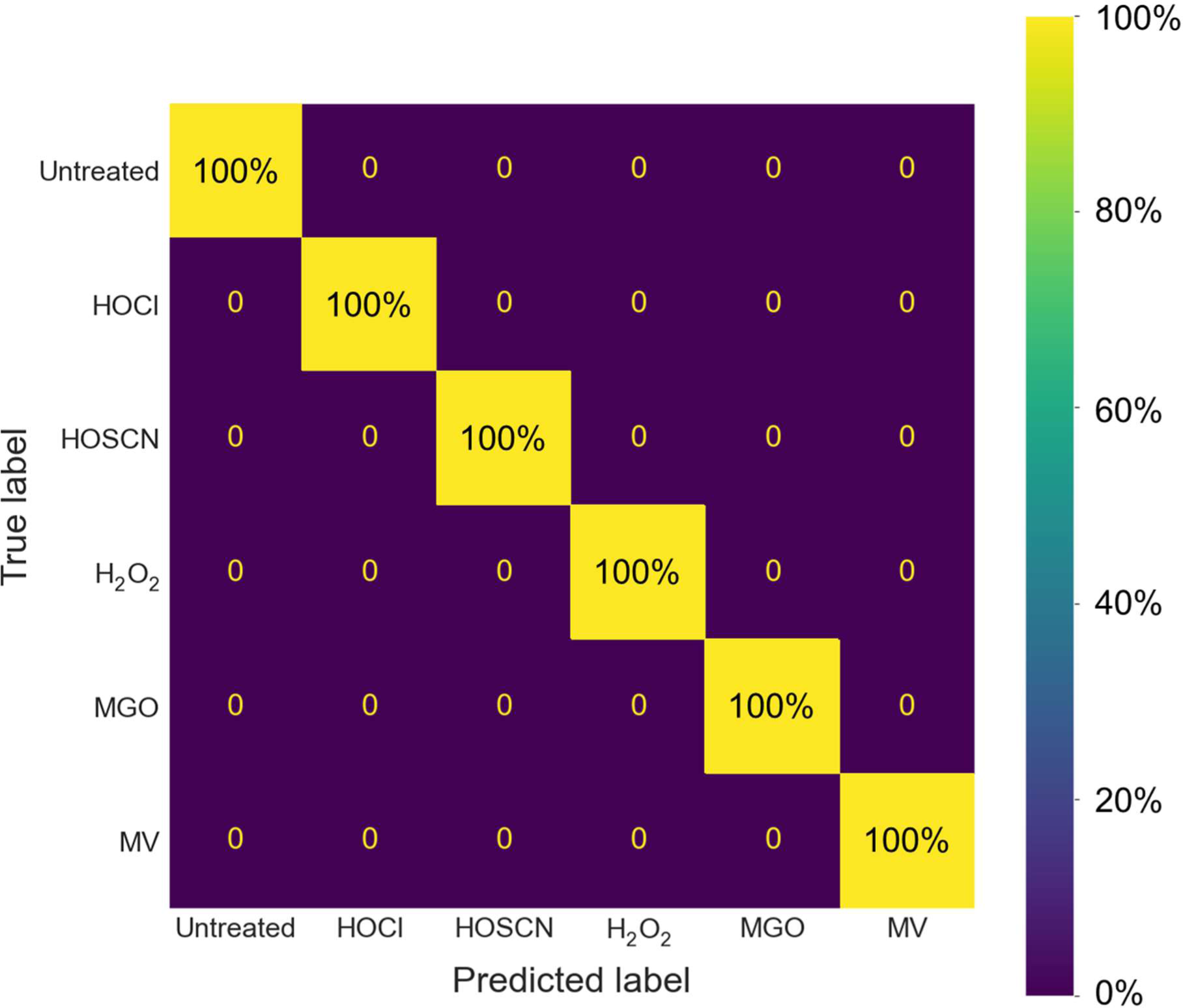
Confusion matrix for the SVM classification of P. aeruginosa samples exposed to oxidative stress conditions. The whole m/z range mass spectra acquired using LD-REIMS was employed to classify each sample into one of six classes. The model was validated using k-fold cross validation (k=10). The samples were classified with 100% accuracy.

### 4. REIMS analysis can identify biomarkers of general oxidative stress and biomarkers specific to individual oxidants

We next investigated whether the metabolic changes detected in *P. aeruginosa* were specific to one or common to two or more oxidative stresses. Features were selected based on the relative importance of their linear regression coefficients obtained during SVM model training. Fifty-four unique metabolites with concentrations significantly impacted by exposure to at least one oxidant were identified (Table S2). Thirty-four of these were previously identified as *P. aeruginosa* metabolites using the same mass spectrometer (Bardin et al., 2018). Twenty-four had their identity confirmed using LC-MS/MS (Table S3).

The changes in metabolite concentration observed in oxidatively stressed samples and the impact of each oxidant on the three classes of biomolecule are shown in Fig. 6 (see Table S4 for a list of the fold-changes in intensity and associated *p* values for each metabolite). These data allow us to draw conclusions about the specific metabolic impact of each oxidant. Exposure to HOCl significantly decreased the concentration of ten AHQs (Fig. 6, Fig. 7) to an extent not seen with any other oxidant, while also causing a decrease the concentration of three rhamnolipids (Fig. 6, Fig. 8). In contrast 14 of the 21 phospholipids impacted by HOCl stress showed an increase in concentration, with the most notable exception being the phosphatidylglycerols PG(35:0), PG(35:1) and PG(35:2) which were significantly lower in concentration for all oxidants beside MV. No AHQ or rhamnolipid was impacted by exposure to HOSCN. However, 19 phospholipids were impacted by HOSCN stress. The phospholipids PE(30:0), PE(32:1) and PG(32:0) had higher concentrations in the HOSCN-treated samples than any other oxidant. Exposure to H2O2 had the strongest impact on rhamnolipid concentrations of all the oxidants (Fig 8), while the impact on phospholipids was similar to the other treatments besides MV. The metabolite profiles of MGO-treated samples were broadly similar to those of H2O2-treated samples, having an almost identical phospholipid profile and having the second strongest impact on the detected rhamnolipids after H2O2. MV had a unique profile compared to the other treatments in the phospholipid region, causing an increase in concentration of several phospholipids that decreased in concentration for all other oxidants, and vice versa. The phospholipids PE(33:2), PE(33:1) and PE(30:0) increased in concentration only upon exposure to MV. MV also had a distinct impact on the growth of *P. aeruginosa* cultures, causing a reduction in the final OD600 reached in stationary phase compared to untreated samples (Fig. S1). Each oxidant except HOSCN elicited at least one oxidant-specific metabolic change.

**Figure 6:**
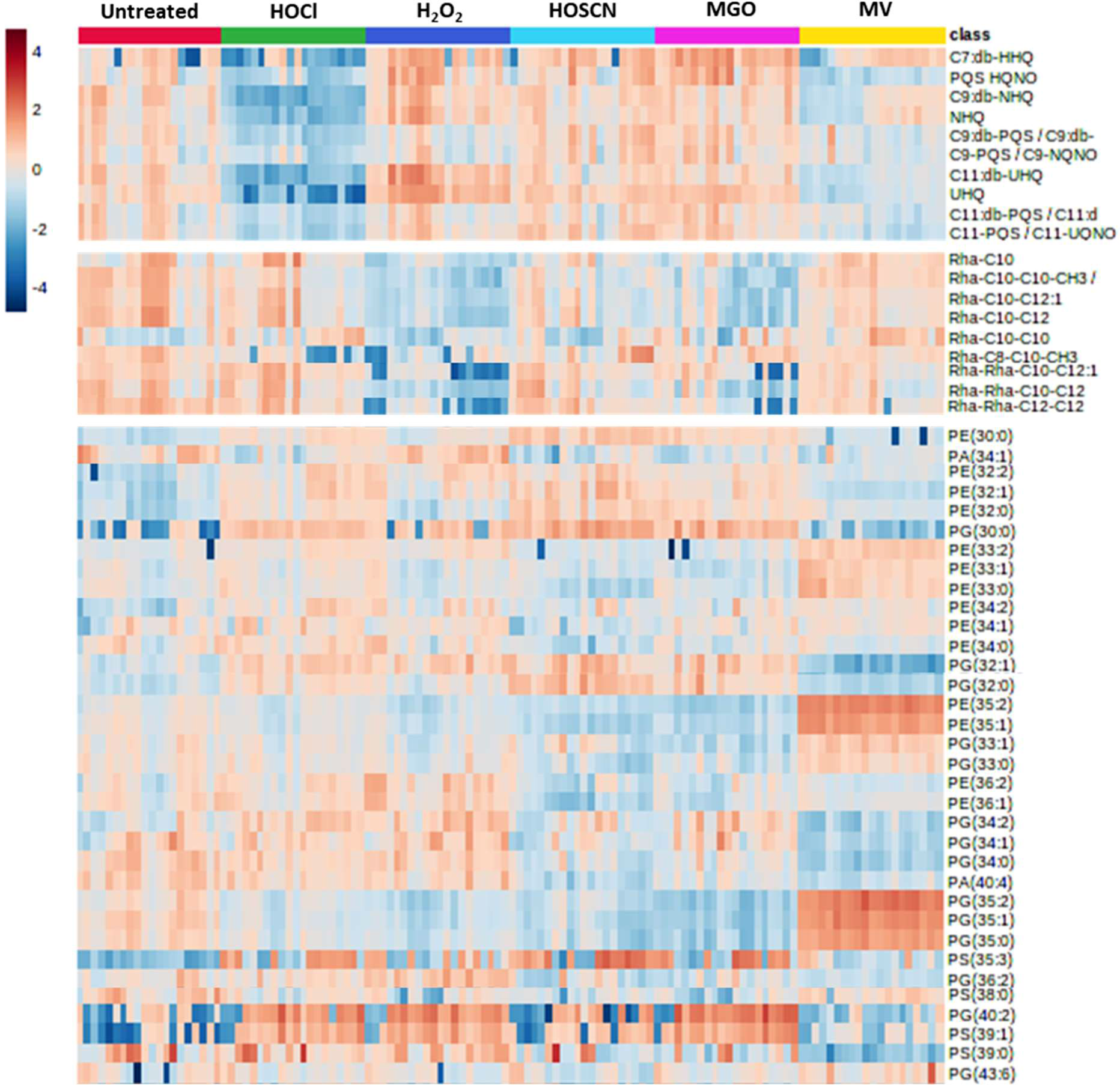
Heatmap visualisation of the results of ANOVA of the metabolite profiles of 10 biological replicates P. aeruginosa grown untreated or in the presence of different oxidative stresses. Colour represents the variation in the intensity of metabolite peaks in mass spectra, relative to the mean, where dark red means a higher intensity and blue a lower intensity. Metabolites are arranged into the three detected classes; PQS associated AHQs, rhamnolipids and phospholipids.

**Figure 7:**
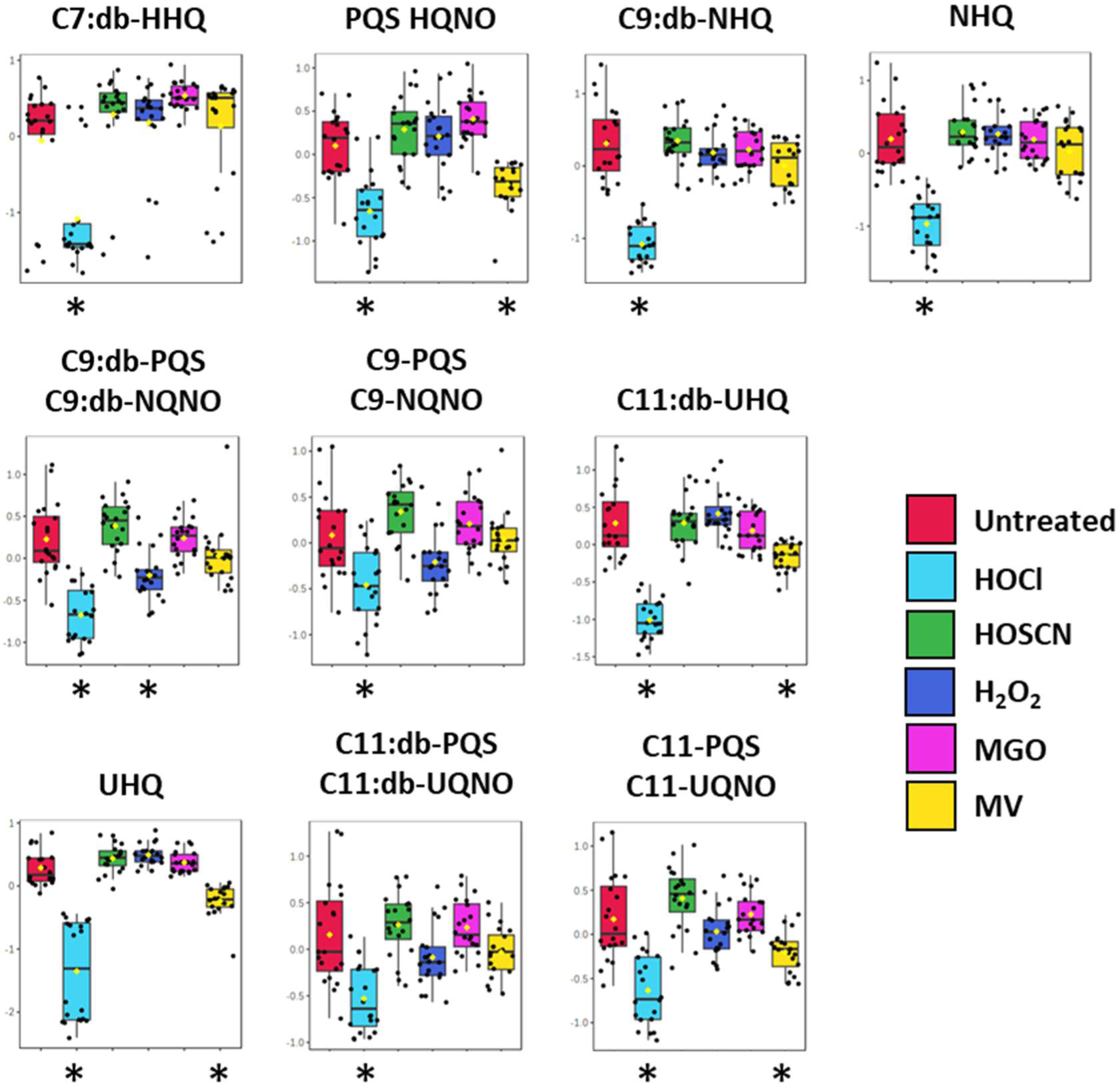
Exposure to HOCl causes a significant decrease in concentration of all 10 PQS-associated quorum sensing molecules identified using LD-REIMS, including the Pseudomonas Quinolone Signal (PQS) and its immediate precursor HHQ, the two most active signalling molecules for the PQS system. MV causes a decrease in the concentration of 4, and H2O2 in 1. The threshold criterion for significance was a p value < 0.05, determined using ANOVA and Fisher’s LSD post-hoc analysis. Classes significantly different from untreated are highlighted with an asterisk. The box and whisker plots summarise the normalised intensity values (mean +/-SD) for the metabolite.

**Figure 8:**
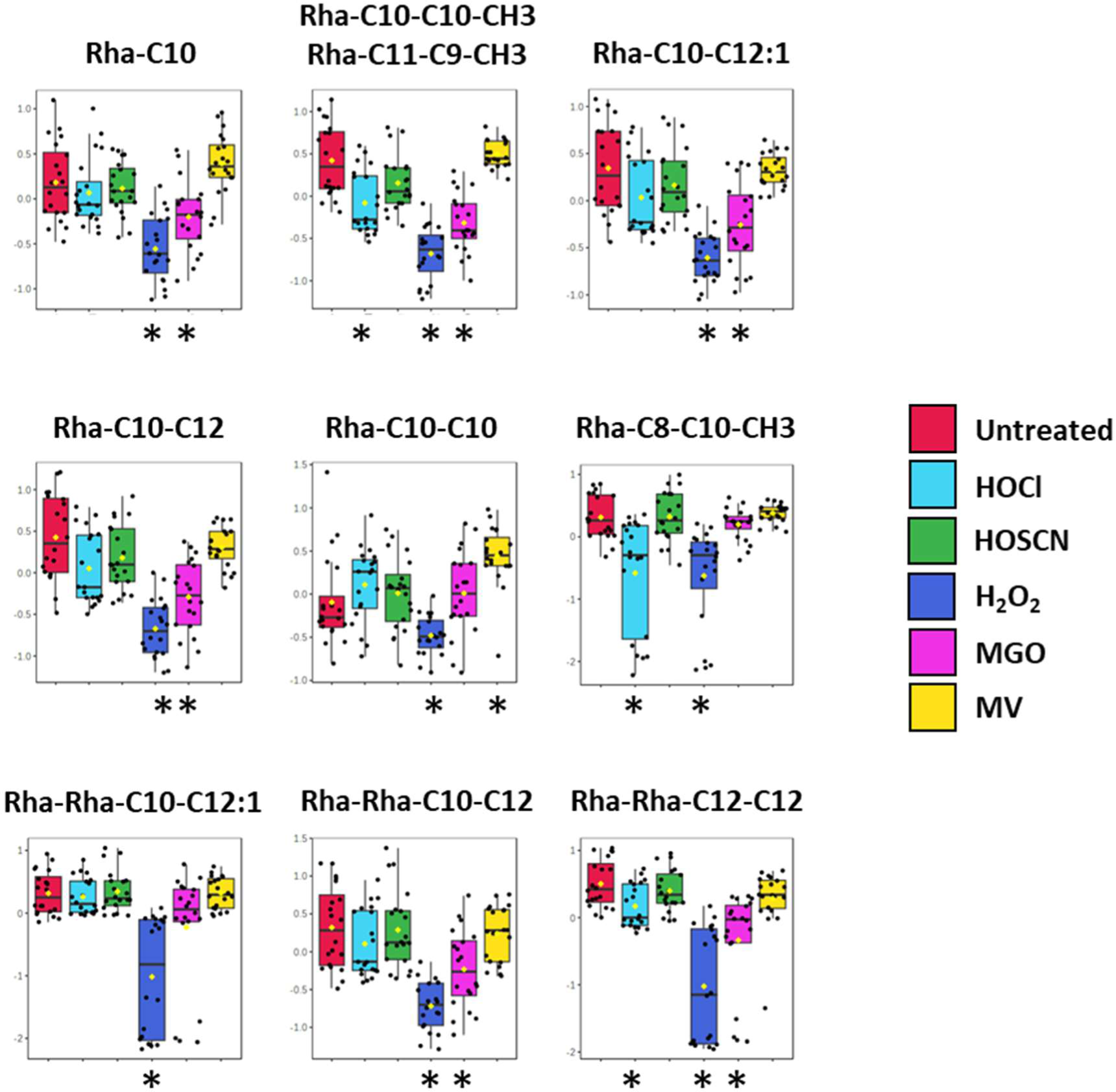
Exposure to H2O2 causes a decrease in concentration of all 9 detected rhamnolipids. MGO causes a decrease in concentration of 5, HOCl in 3 and MV in 1. The threshold criterion for significance was a p value < 0.05, determined using ANOVA and Fisher’s LSD post-hoc analysis. Classes significantly different from untreated are highlighted with an asterisk. The box and whisker plots summarise the normalised intensity values (mean +/-SD) for the metabolite.

### 5. Biomarkers of HOCl and HOSCN stress are conserved in *P. aeruginosa* clinical isolates

Our data has established that LD-REIMS can discriminate between oxidatively stressed and untreated and identified biomarkers specific to exposure to different oxidants in PA14 samples, a commonly used lab strain of *P. aeruginosa*. For these findings to have potential use in the investigation of *P. aeruginosa* infections *in vivo*, for example to establish whether bacteria are exposed to these stresses during infection, we needed to demonstrate that these specific metabolomic responses are conserved in clinical strains. Therefore, we carried out LD-REIMS analysis on 10 *P. aeruginosa* strains isolated from Cystic Fibrosis (CF) lung infections and compared their untreated and HOCl-treated metabolomes using LD-REIMS.

Separation in PCA scores plots between treated and untreated samples was reduced compared to the lab strain analysis (Fig. S4, Fig. S5). However, this included the PA14 control, suggesting the reduced discriminative power was due to a lower number of replicates (n = 6 compared to n = 20).

We then tracked in the clinical isolates the biomarkers of HOCl and HOSCN established using PA14. The decrease in concentration of AHQs was conserved in the HOCl-treated clinical isolates (Table S5). C9:db-NHQ, C11:db-UHQ and UHQ were conserved in all or all but one isolate, suggesting any impact of HOCl on the PQS system is not limited to the laboratory strain. The rhamnolipids and phospholipid biomarkers were poorly conserved in both the clinical isolates and the PA14 control. The HOSCN biomarkers were very well conserved (Table S6). All biomarkers besides PA(34:1) and PS(38:0) were conserved in the majority of clinical isolates. Only C13 and C25 presented with fewer than 50 % of the biomarkers.

### 6. HOCl-stress reduces quorum sensing-mediated virulence factor levels

The decrease in concentration of ten AHQs associated with the PQS quorum sensing system on exposure to HOCl was one of the strongest patterns observed in our data. While it is not possible to say whether the decrease in concentration is due to a reduction in their production or to direct oxidation of the metabolites, the outcome is likely to be a decrease in the efficacy of the molecules to modulate the PQS-mediated quorum sensing response of *P. aeruginosa*. To investigate this, we measured the concentration of pyocyanin and elastase, virulence factors known to be regulated by the PQS system, in WT cultures grown in HOCl. We compared the levels to those in *pgsE* mutants, which are mutated in genes required for PQS synthesis and are expected to produce little or no pyocyanin or elastase (Bala et al., 2015). Pyocyanin and elastase were clearly made by the wild-type strain but when grown in the presence of HOCl the levels were significantly reduced (Fig. 9). To test whether the decrease in pyocyanin was due to a decrease in production or direct oxidation, we then treated stationary phase cultures in which pyocyanin was already present with HOCl and measured the changing concentration over time. We did not see a difference in the concentration of pyocyanin in treated cultures (Fig. S8), suggesting that HOCl does not cause direct oxidation of pyocyanin.

**Figure 9:**
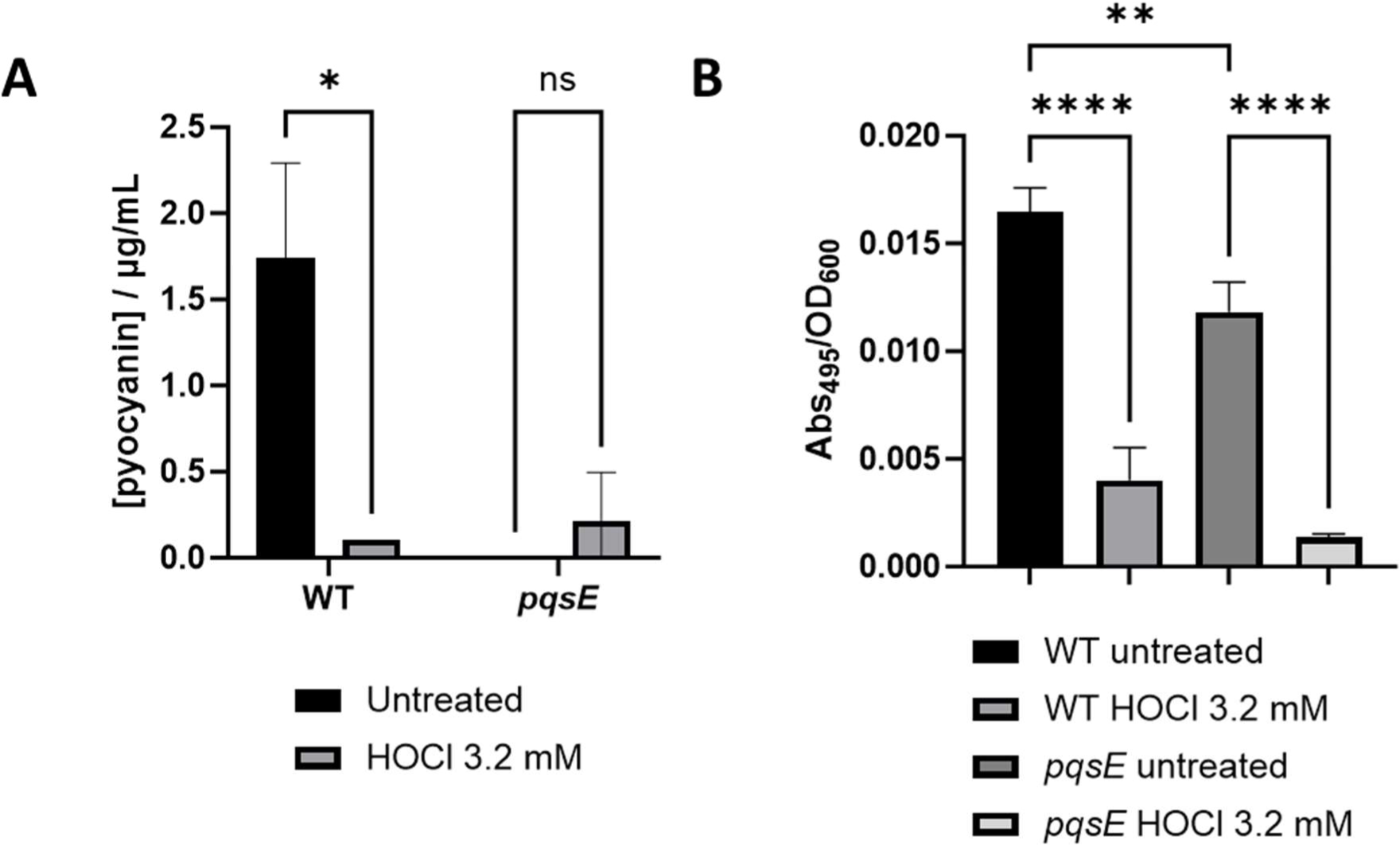
P. aeruginosa cultures grown in the presence of HOCl (3.2 mM) are deficient in pyocyanin (A) and elastase (B) compared to untreated.

## DISCUSSION

### LD-REIMS can be used to confidently distinguish between oxidatively stressed *P. aeruginosa* cultures

In this study, we tested LD-REIMS as a novel technique for the metabolic profiling of *P. aeruginosa*, with the aim to identify biomarkers of oxidative stress and to provide insight into the oxidative stress protection systems employed by the bacteria. REIMS has been employed for a range of microbiological studies, most frequently for bacterial identification and classification (Bodai et al., 2018; Bolt et al., 2016; Cameron et al., 2016, 2021; Strittmatter et al., 2014), as well as metabolic screening of yeast colonies (Gowers et al., 2019) and imaging (Golf et al., 2015). A study in the Takats’ lab successfully classified *P. aeruginosa* isolates to a strain level with 81 % accuracy and identified a number of *P. aeruginosa* metabolites (Bardin et al., 2018).

Whilst REIMS has been established for the analysis of bacterial colonies, the LD-REIMS protocol reported here allows for the analysis of pelleted samples, meaning bacterial samples grown in liquid, including high salt media that REIMS would normally be intolerant towards, can be analysed in a high-throughput manner. We analysed and compared the metabolomes of *P. aeruginosa* grown in the absence and presence of oxidative stresses, with the only sample preparation requirement being the centrifugation and washing of the samples just prior to analysis. Using our approach, 96 samples can be analysed in 20 minutes, and samples are analysed within 60 minutes of the start of sample preparation. Additionally, this was the first reported application of a 3 μm laser for high throughout REIMS, after ongoing research found that a 3 μm, shorter pulse width laser led to increased ion yields compared to the original CO2 setup (Simon et al., 2023).

The limitation of REIMS is that it has a restricted metabolic coverage, largely detecting hydrophobic molecules in the surface layers of bacteria, but molecules that will be first to come into contact with an externally applied stress.

We used the LD-REIMS-derived mass spectra to classify *P. aeruginosa* PA14 samples using a support vector machine classification model based on the specific oxidative stress they were grown in with 100% accuracy. This result was reproducible, with classification accuracy consistent for samples prepared and analysed on different days.

The lab strain PA14 is a burn wound isolate that represents the most common clonal group worldwide and is more virulent than other commonly used strains PAO1 and PAK (Grace et al., 2022). It is now recognised that it is important not to rely on analysis of a limited number of laboratory reference strains but to extend the natural isolates, including clinical strains (Hefnawy et al., 2018). For this reason, we extended our LD-REIMS study to clinical isolates from CF patients. The biomarkers of HOCl and HOSCN stress were conserved in the clinical isolates.

### A unique metabolic profile was recorded for each oxidant

The observed classification accuracy of 100% indicates that growth in the presence of different oxidants impacts the *P. aeruginosa* metabolome in a manner unique to each oxidant, at least for the metabolites detectable using LD-REIMS. We were able to use a combination of univariate and multivariate analyses to find the metabolites with significantly different intensities between sample classes. These included representatives from all three of the main classes of *P. aeruginosa* metabolites detected using LD-REIMS: AHQs associated with the PQS quorum sensing system, rhamnolipids and phospholipids (Bardin et al., 2018).

Of the oxidants, HOCl and MV (superoxide) had the greatest impact on the *P. aeruginosa* metabolome, both seen to cluster away from the untreated samples in PLS-DA. HOSCN, H2O2 and MGO did not separate from untreated when all oxidants were analysed together, however clear separation from untreated samples was observed when analysed independently using two-group analyses like OPLS-DA, and samples were classified with 100% accuracy. This suggests HOCl and MV have a broader impact on the metabolome, disrupting the concentrations of a greater range of molecules, while the other three impact fewer metabolites but still leave a consistent and unique mark on the metabolome. The greater impact of HOCl matches with the reaction profiles of the oxidants. HOCl is a stronger oxidant than H2O2 or HOSCN, able to react with a wide range of bacterial substrates including thiols and amines (Winterbourn et al., 2016). HOSCN reacts almost solely with thiol groups, while H2O2 only reacts directly with thiols and transition metal centres, due to the high activation energy of its reactions (Winterbourn & Hampton, 2008). Reaction profiles are not sufficient to explain the greater impact of MV, as superoxide has a low reactivity in aqueous solutions (Halliwell & Gutteridge, 2015). However, the use of methyl viologen as an intermediary for superoxide generation may explain the impact on the metabolome. MV (also known as paraquat) is non-polar and can therefore pass through bacterial membranes, where superoxide is generated within the cell. The intracellular production of MV would likely have a significantly different impact on the metabolome compared to the other oxidants, which may cause more damage at the cell membrane. This is particularly relevant for this study, as the detected molecules are membrane-associated or extracellular.

### HOCl has a specific impact on AHQs associated with the PQS quorum sensing system

Our most striking finding was the decrease in concentration of ten AHQs associated with the PQS quorum sensing system after exposure to HOCl. Among these are 2-heptyl-3-hydroxy-4(1*H*)-quinolone, also known as the *Pseudomonas* quinolone signal (PQS), the most active signal molecule of the Pqs system that regulates the production of virulence factors including elastase, rhamnolipids, pyocyanin and siderophores such as pyoverdine, alongside inducing its own production (García-Reyes et al., 2020).

The decrease in concentration of AHQs was also seen in the clinical isolates (Table S6), clearly showing that the impact of HOCl on the PQS system is replicated in clinical strains.

PQS and other AHQs are released extracellularly by *P. aeruginosa* where their concentration acts as a proxy of the local bacterial population and subsequently dictates the expression of PQS-dependent genes. Due to their alkyl chain moiety, AHQs are hydrophobic and rely on membrane vesicles (MVs) for transport to and excretion from the cell. Their association with MVs and the cell envelope explains their prevalence in LD-REIMS *P. aeruginosa* mass spectra alongside phospholipids and rhamnolipids. The PQS system is associated with the oxidative stress response of *P. aeruginosa*, having been shown to act as both an antioxidant and pro-oxidant in different contexts (Bollinger et al., 2001; García-Reyes et al., 2020; Häussler & Becker, 2008). Its antioxidant effect is attributed its nature as a strong electron donor due to its bisoxygenated aromatic structure and can therefore reduce intracellular reactive oxygen species (ROS) levels through direct reaction with oxidants. However, it has also been shown to sensitise *P. aeruginosa* to killing by H2O2 and oxidative stress mediated via ultraviolet radiation (Häussler & Becker, 2008; Pezzoni et al., 2015). This pro-oxidant effect is thought to be due to the ability of PQS to chelate Fe^3+^ ions, resulting in the formation of free radicals.

It is unclear whether the decrease in concentration of the AHQs is due to HOCl inhibiting their production or to their direct oxidation. However, irrespective of the mechanism, the phenotypic impact is clear as the concentrations of PQS-regulated virulence factors pyocyanin and elastase were reduced in HOCl stressed cultures to the levels seen in PQS-deficient mutant strains. In addition, we showed that this was not due to oxidation of pyocyanin by HOCl, supporting a model whereby HOCl prevents the production of AHQs. Alongside prevention of virulence factor production, this would additionally remove the potential antioxidant properties of the PQS system. These findings lead to the hypothesis that neutrophil generated HOCl formation in response to infection, whether it is generated following phagocytosis or in neutrophil extracellular traps following NETosis, could result in reduced PQS-mediated QS and virulence factor formation in the infecting *P. aeruginosa* population, augmenting the direct impact of neutrophil killing mechanisms.

## SUPPLEMENTARY

**Figure S1:**
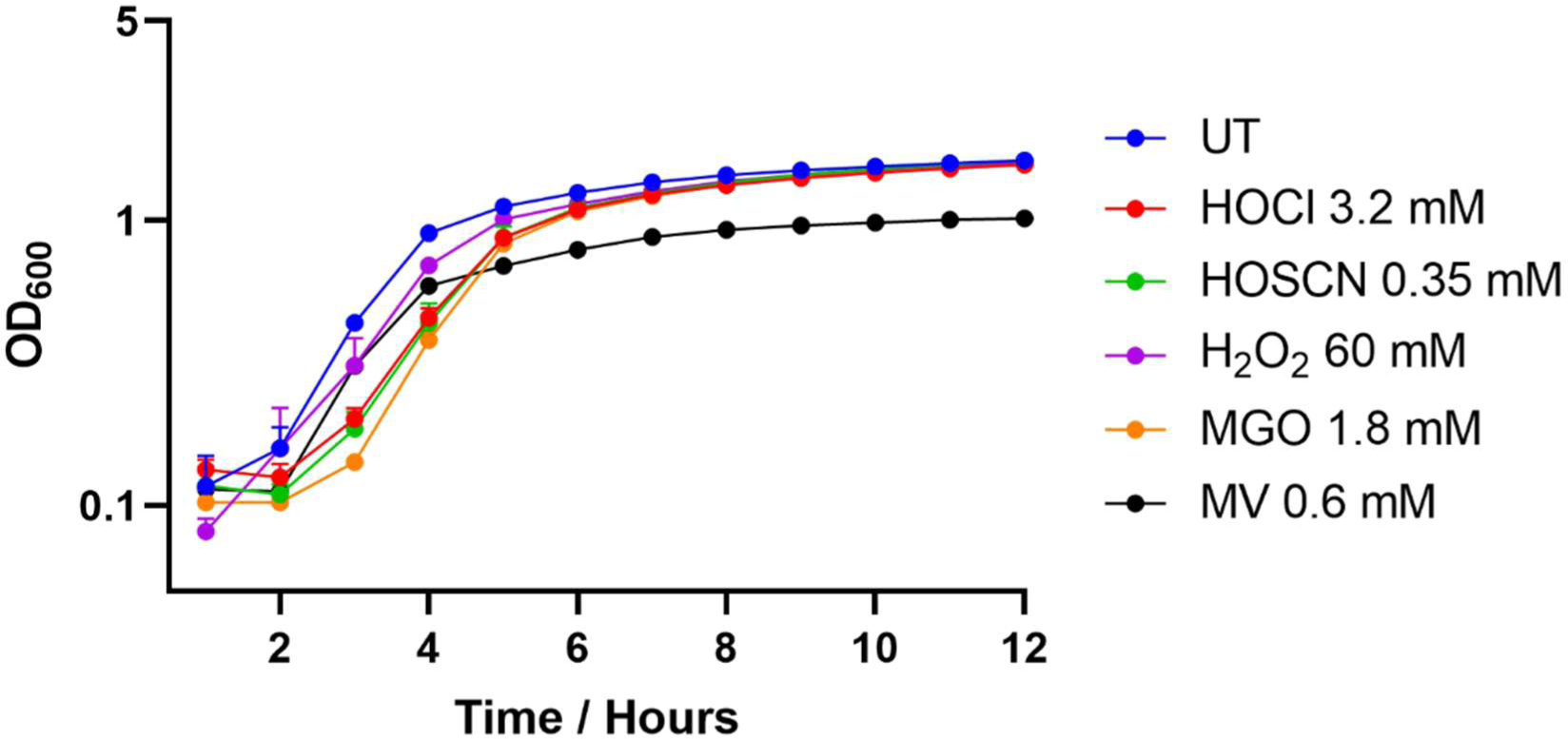
Growth data for untreated samples compared to samples treated with oxidants at the concentrations employed in this study. Bacteria were subcultured into untreated or treated LB in a 96-well plate format and incubated overnight at 37 °C, shaking at 700 rpm. Points represent the mean OD600 value at each timepoint, whilst the error bars represent the standard deviation of the mean, (n=6). A range of concentrations were tested, with the aim to find a concentration for each oxidant that provided a one-hour lag in growth compared to untreated. This criterion was chosen to ensure the bacteria were responding to the stress whilst not being killed.

**Figure S2:**
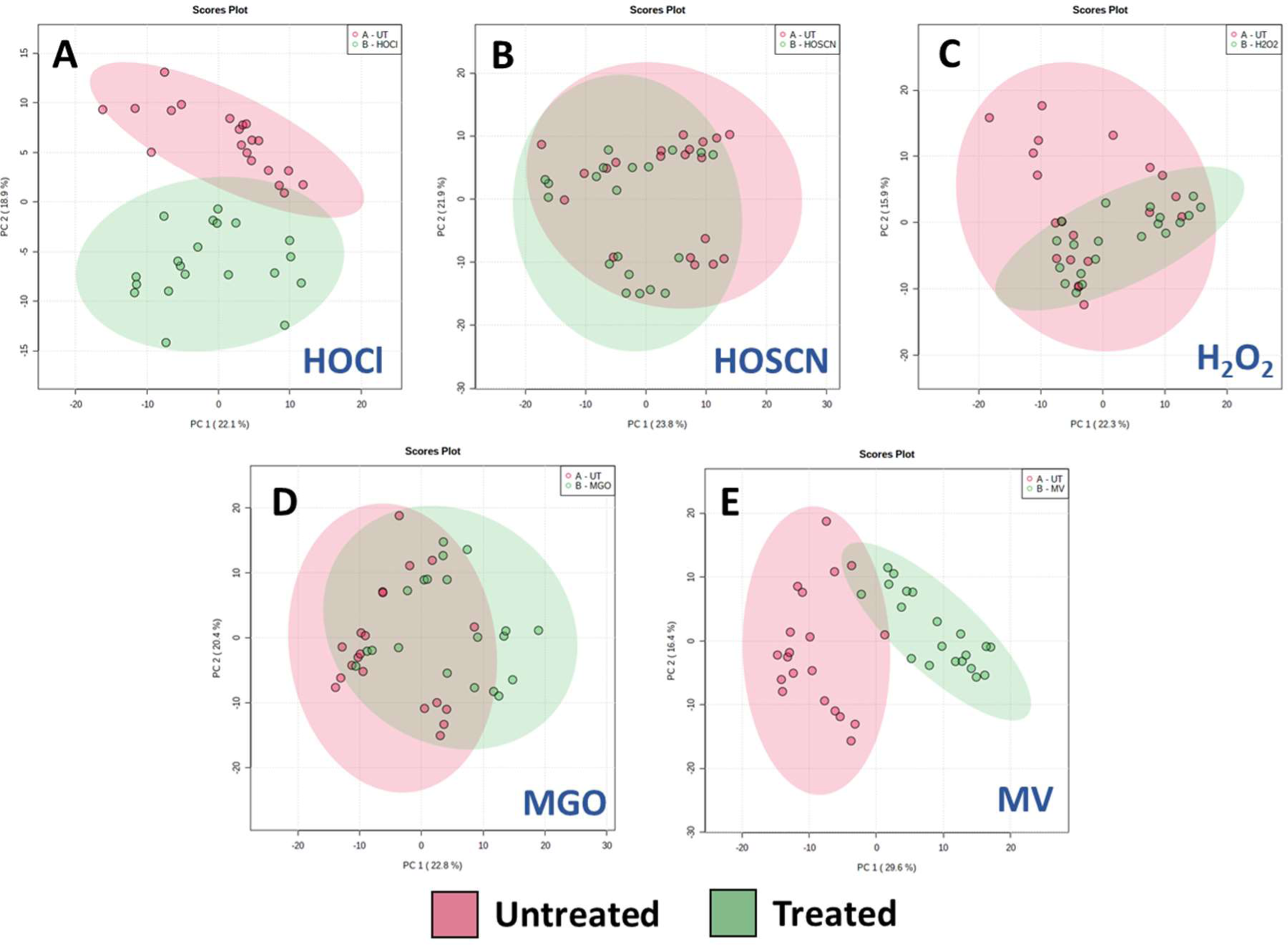
Statistical analysis of P. aeruginosa mass spectrum data obtained by LD-REIMS by a two-group PCA of untreated and treated P. aeruginosa samples; (a) 3.2 mM HOCl, (b) 0.35 mM HOSCN, (c) 60 mM H2O2; (d) 1.8 mM MGO (e) 0.6 mM MV. Samples were prepared and analysed over two independent days, n=20 (10 per day). Statistical analyses were completed on the 50 to 1200 m/z range, after background subtraction and mass drift correction, using the MetaboAnalyst 5.0 platform. The plots all represent components one and two. Shaded areas show 95% confidence intervals of the sample groups. The HOCl- and MV-treated samples (panels A and E respectively) clearly separate from the untreated, suggesting a strong impact of the metabolic profile. The samples for the other treatments (HOSCN, H2O2 and MGO, panels B, C and E respectively) do not clearly separate from untreated at the concentration used.

**Figure S3:**
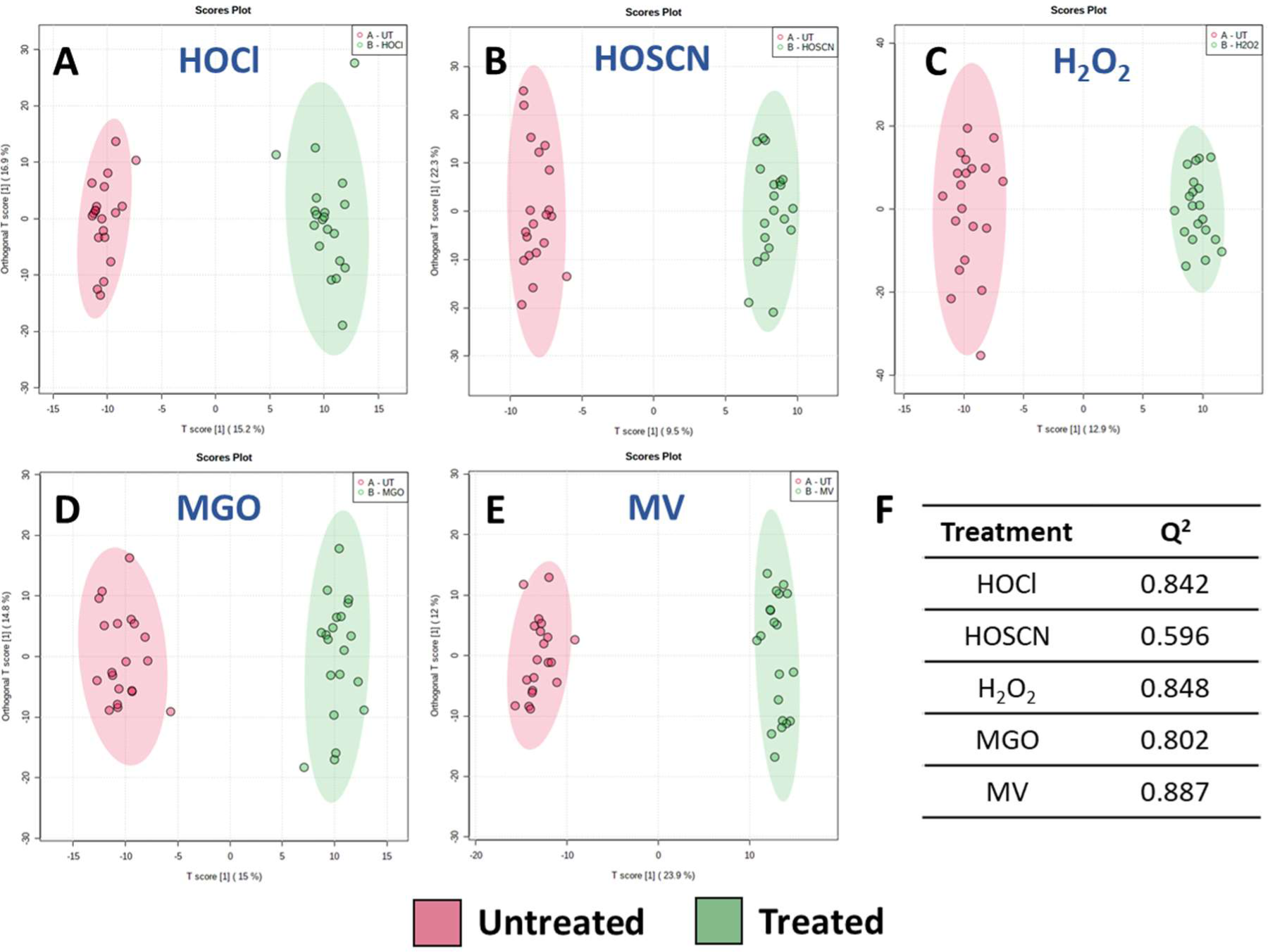
Statistical analysis of P. aeruginosa mass spectrum data obtained by LD-REIMS by OPLS-DA analysis. Score plots generated based on OPLS-DA of untreated and treated P. aeruginosa samples; (a) 3.2 mM HOCl, (b) 0.35 mM HOSCN, (c) 60 mM H2O2; (d) 1.8 mM MGO (e) 0.6 mM MV. Samples were prepared and analysed over two independent days, n=20 (10 per day). Statistical analyses were completed on the 50 to 1200 m/z range, after background subtraction and mass drift correction, using the MetaboAnalyst 5.0 platform. The model is able to separate the untreated and treated samples for each oxidant, cross-validated by the high Q2 coefficient for each plot (see table).

**Figure S4:**
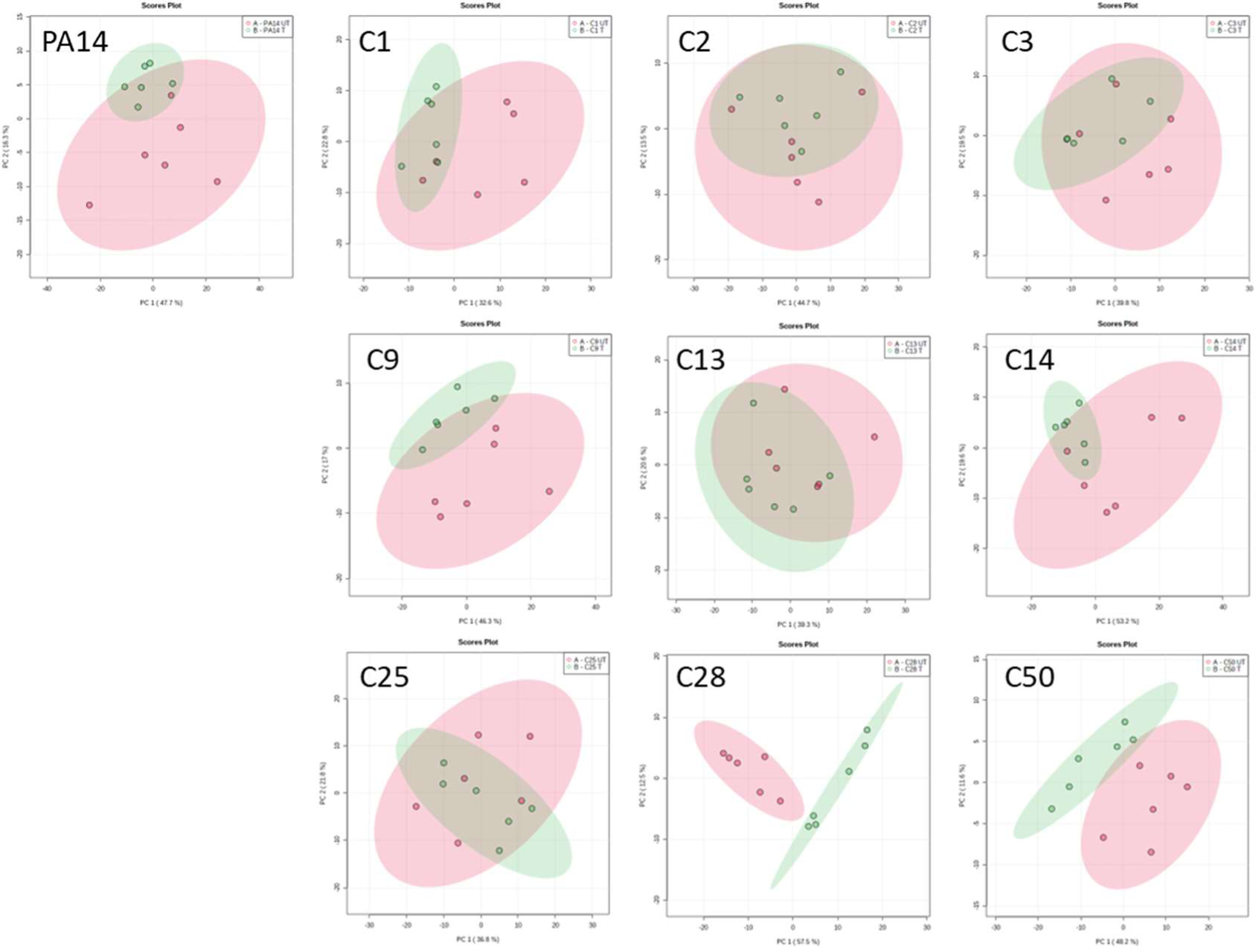
Two-group PCA of untreated and HOCl-treated P. aeruginosa strains, comparing PA14 to 10 cystic fibrosis (CF) isolates (n=6). Statistical analyses were completed on the 50 to 1200 m/z range, after background subtraction and mass drift correction, using the MetaboAnalyst 5.0 platform. Separation between untreated and treated samples to the same degree as PA14 is apparent for C1, C9, C13, C14, C15, C28 and C50, suggesting the response to HOCl is conserved in these strains.

**Figure S5:**
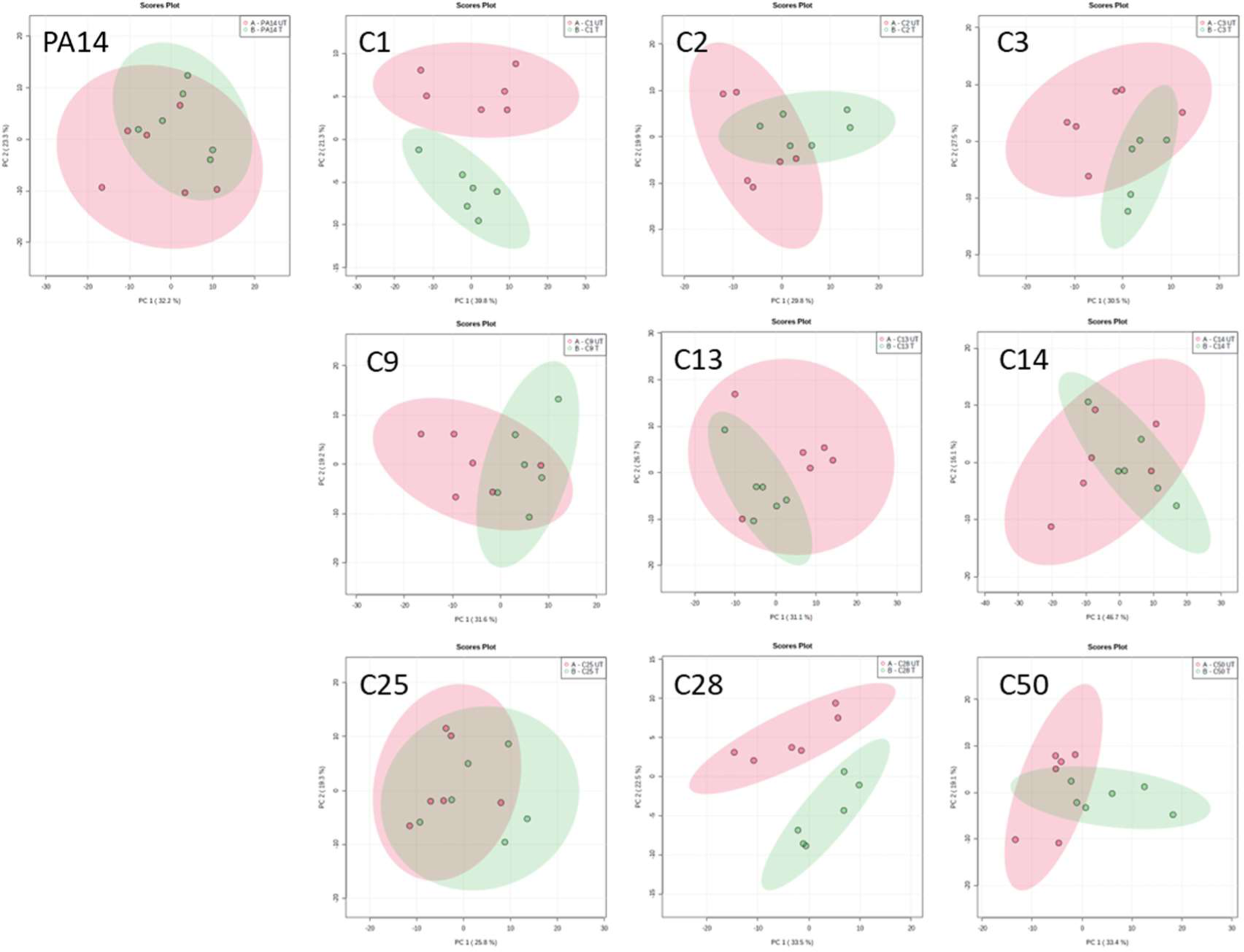
Two-group PCA of untreated and HOSCN-treated P. aeruginosa strains, comparing PA14 to 10 cystic fibrosis (CF) isolates (n=6). Statistical analyses were completed on the 50 to 1200 m/z range, after background subtraction and mass drift correction, using the MetaboAnalyst 5.0 platform. Separation between untreated and treated samples to the same degree as PA14

**Figure S8:**
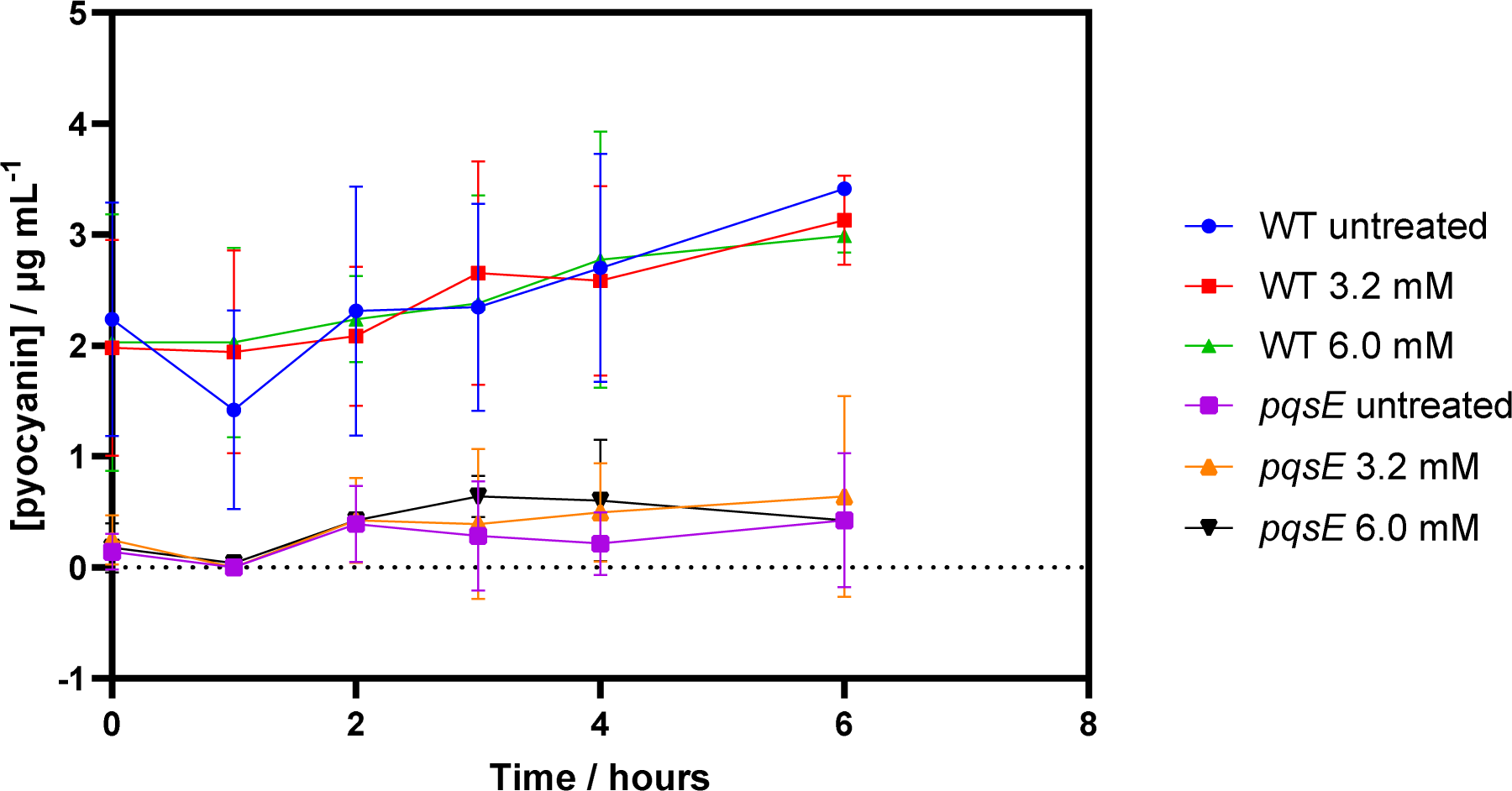
Pyocyanin concentrations in stationary phase P. aeruginosa cultures were not impacted by treatment with HOCl, suggesting the decrease in concentration seen in growing cultures exposed to HOCl is not due to direct oxidisation of pyocyanin.

**Table S1:** P. aeruginosa strains used in this study.

| Strain | Description | Source |
| --- | --- | --- |
| PA14 (WT) | Wild-type lab stock | M. Ausubel, Harvard (Rahme et al., 1995) |
| *pqsE* | PAMr: PA14 wild-type background, MAR2xT7 transposon, Gm<sup>r</sup>. PqsE Quinolone signal response protein. | M. Ausubel, Harvard (Liberati NT et al., 2006) |
| C1 | *P. aeruginosa* clinical isolate from early infection of CF lung | Colindale |
| C2 | *P. aeruginosa* clinical isolate from early infection of CF lung | Colindale |
| C3 | *P. aeruginosa* clinical isolate from early infection of CF lung | Colindale |
| C9 | *P. aeruginosa* clinical isolate from early infection of CF lung | Colindale |
| C28 | *P. aeruginosa* clinical isolate from early infection of CF lung | Colindale |
| C50 | *P. aeruginosa* clinical isolate from early infection of CF lung | Colindale |
| C12 | *P. aeruginosa* clinical isolate from chronic infection of CF lung | Royal Brompton Hospital |
| C13 | *P. aeruginosa* clinical isolate from chronic infection of CF lung | Royal Brompton Hospital |
| C14 | *P. aeruginosa* clinical isolate from chronic infection of CF lung | Royal Brompton Hospital |
| C15 | *P. aeruginosa* clinical isolate from chronic infection of CF lung | Royal Brompton Hospital |
| C17 | *P. aeruginosa* clinical isolate from chronic infection of CF lung | Royal Brompton Hospital |
| C25 | *P. aeruginosa* clinical isolate from chronic infection of CF lung | Royal Brompton Hospital |

**Table S2:** All metabolites significantly impacted by oxidant treatment identified to at least MSI3.

| Annotation | Detected ion | Accurate m/z | Theoretical m/z | Error (ppm) | MSI |
| --- | --- | --- | --- | --- | --- |
| C7:db-HHQ | [M-H]- | 240.1386 | 240.1394 | -3.331 | 3 |
| PQS HQNO | [M-H]- | 258.1493 | 258.1500 | -2.712 | 3 |
| C9:db-NHQ | [M-H]- | 268.1699 | 268.1707 | -2.983 | 3 |
| NHQ | [M-H]- | 270.1857 | 270.1863 | -2.221 | 2 |
| C9:db-PQS<br>C9:db-NQNO | [M-H]- | 284.1648 | 284.1656 | -2.815 | 3 |
| C9-PQS<br>C9-NQNO | [M-H]- | 286.1809 | 286.1813 | -1.398 | 3 |
| C11:db-UHQ | [M-H]- | 296.2010 | 296.2020 | -3.376 | 2 |
| UHQ | [M-H]- | 298.2173 | 298.2176 | -1.006 | 2 |
| C11:db-PQS<br>C11:db-UQNO | [M-H]- | 312.1958 | 312.1969 | -3.523 | 2 |
| C11-PQS<br>C11-UQNO | [M-H]- | 314.2105 | 314.2126 | -6.683 | 2 |
| Rha-C10 | [M-H]- | 333.1909 | 333.1914 | -1.501 | 3 |
| Rha-C10-C10-CH3<br>Rha-C11-C9-CH3 | [M-H]- | 517.3356 | 517.3377 | -4.059 | 3 |
| Rha-C10-C12:1 | [M-H]- | 529.3377 | 529.3377 | 0.000 | 3 |
| Rha-C10-C12 | [M-H]- | 531.3534 | 531.3534 | 0.000 | 2 |
| Rha-C10-C10 | [M+Cl]- | 539.2986 | 539.2987 | -0.185 | 3 |
| Rha-C8-C10-CH3 | [M-H]- | 545.3596 | 545.3690 | -17.236 | 3 |
| Rha-Rha-C10-C12:1 | [M-H]- | 675.3931 | 675.3956 | -3.702 | 3 |
| Rha-Rha-C10-C12 | [M-H]- | 677.4106 | 677.4113 | -1.033 | 2 |
| Rha-Rha-C12-C12 | [M-H]- | 705.4439 | 705.4431 | 1.134 | 3 |
| PE(30:0) | [M-H]- | 662.4751 | 662.4761 | -1.509 | 2 |
| PA(34:1) | [M-H]- | 673.4812 | 673.4814 | -0.297 | 3 |
| PE(32:2) | [M-H]- | 686.4746 | 686.4766 | -2.913 | 3 |
| PE(32:1) | [M-H]- | 688.4902 | 688.4923 | -3.050 | 2 |
| PE(32:0) | [M-H]- | 690.5051 | 690.5079 | -4.055 | 2 |
| PG(30:0) | [M-H]- | 693.4734 | 693.4707 | 3.893 | 2 |
| PE(33:2) | [M-H]- | 700.4902 | 700.4918 | -2.284 | 3 |
| PE(33:1) | [M-H]- | 702.5056 | 702.5074 | -2.562 | 2 |
| PE(33:0) | [M-H]- | 704.5186 | 704.5231 | -6.387 | 2 |
| PE(34:2) | [M-H]- | 714.5062 | 714.5079 | -2.379 | 3 |
| PE(34:1) | [M-H]- | 716.5231 | 716.5231 | 0.000 | 2 |
| PE(34:0) | [M-H]- | 718.5305 | 718.5387 | -11.412 | 3 |
| PG(32:1) | [M-H]- | 719.4958 | 719.4868 | 12.509 | 2 |
| PG(32:0) | [M-H]- | 721.4997 | 721.5020 | -3.188 | 2 |
| PE(35:2) | [M-H]- | 728.5221 | 728.5236 | -2.059 | 3 |
| PE(35:1) | [M-H]- | 730.5365 | 730.5392 | -3.696 | 2 |
| PG(33:1) | [M-H]- | 733.5014 | 733.5020 | -0.818 | 2 |
| PG(33:0) | [M-H]- | 735.5128 | 735.5177 | -6.662 | 2 |
| PE(36:2) | [M-H]- | 742.5399 | 742.5392 | 0.943 | 3 |
| PE(36:1) | [M-H]- | 744.5514 | 744.5544 | -4.029 | 3 |
| PG(34:2) | [M-H]- | 745.5013 | 745.5025 | -1.610 | 2 |
| PG(34:1) | [M-H]- | 747.5173 | 747.5181 | -1.070 | 3 |
| PG(34:0) | [M-H]- | 749.5251 | 749.5244 | 0.934 | 3 |
| PA(40:4) | [M-H]- | 751.5321 | 751.5278 | 5.722 | 3 |
| PG(35:2) | [M-H]- | 759.5166 | 759.5181 | -1.975 | 2 |
| PG(35:1) | [M-H]- | 761.5314 | 761.5333 | -2.495 | 2 |
| PG(35:0) | [M-H]- | 763.5314 | 763.5283 | 4.060 | 2 |
| PS(35:3) | [M-H]- | 770.4961 | 770.4973 | -1.557 | 3 |
| PG(36:2) | [M-H]- | 773.5316 | 773.5338 | -2.844 | 3 |
| PS(37:3) | [M-H]- | 798.5255 | 798.5286 | -3.882 | 3 |
| PS(38:0) | [M-H]- | 818.5872 | 818.5912 | -4.886 | 3 |
| PG(40:2) | [M-H]- | 829.5879 | 829.5959 | -9.643 | 3 |
| PS(39:1) | [M-H]- | 830.5886 | 830.5912 | -3.130 | 3 |
| PS(39:0) | [M-H]- | 832.6060 | 832.6068 | -0.961 | 3 |
| PG(43:6) | [M-H]- | 863.5857 | 863.5830 | 3.127 | 3 |

**Table S3:**
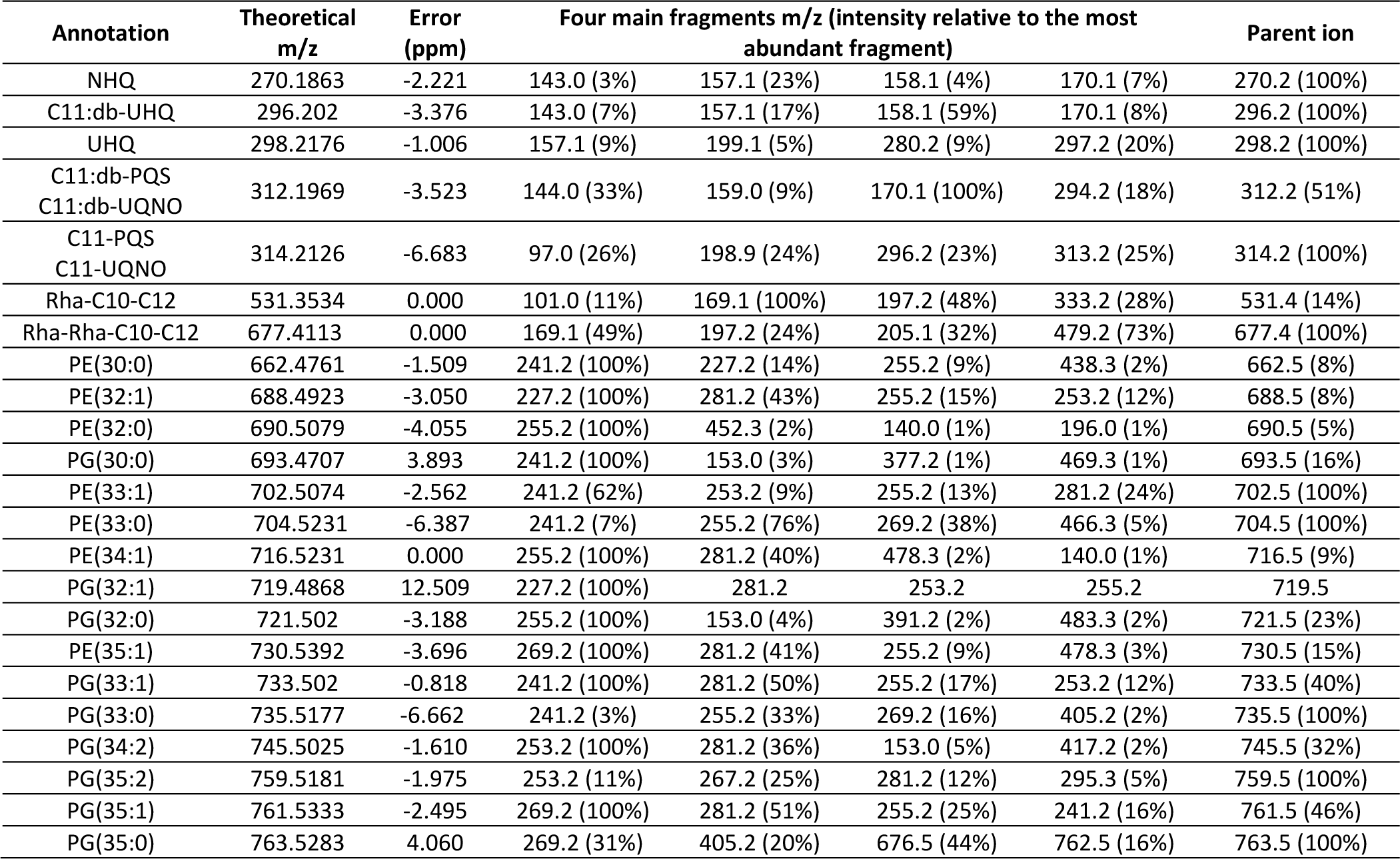
Identification of 24 of the most significant molecules based on MSMS spectral data acquired with LCMS analysis. The four most intense fragment ions are given together their parent ion exact mass. Intensities are given related to the most intense fragment ion. Unattributed fragment ions were included in the table; isotope ions were excluded.

| Annotation | Theoretical m/z | Error (ppm) | Four main fragments m/z (intensity relative to the most abundant fragment) |  |  |  | Parent ion |
| --- | --- | --- | --- | --- | --- | --- | --- |
| NHQ | 270.1863 | -2.221 | 143.0 (3%) | 157.1 (23%) | 158.1 (4%) | 170.1 (7%) | 270.2 (100%) |
| C11:db-UHQ | 296.202 | -3.376 | 143.0 (7%) | 157.1 (17%) | 158.1 (59%) | 170.1 (8%) | 296.2 (100%) |
| UHQ | 298.2176 | -1.006 | 157.1 (9%) | 199.1 (5%) | 280.2 (9%) | 297.2 (20%) | 298.2 (100%) |
| C11:db-PQS<br>C11:db-UQNO | 312.1969 | -3.523 | 144.0 (33%) | 159.0 (9%) | 170.1 (100%) | 294.2 (18%) | 312.2 (51%) |
| C11-PQS<br>C11-UQNO | 314.2126 | -6.683 | 97.0 (26%) | 198.9 (24%) | 296.2 (23%) | 313.2 (25%) | 314.2 (100%) |
| Rha-C10-C12 | 531.3534 | 0.000 | 101.0 (11%) | 169.1 (100%) | 197.2 (48%) | 333.2 (28%) | 531.4 (14%) |
| Rha-Rha-C10-C12 | 677.4113 | 0.000 | 169.1 (49%) | 197.2 (24%) | 205.1 (32%) | 479.2 (73%) | 677.4 (100%) |
| PE(30:0) | 662.4761 | -1.509 | 241.2 (100%) | 227.2 (14%) | 255.2 (9%) | 438.3 (2%) | 662.5 (8%) |
| PE(32:1) | 688.4923 | -3.050 | 227.2 (100%) | 281.2 (43%) | 255.2 (15%) | 253.2 (12%) | 688.5 (8%) |
| PE(32:0) | 690.5079 | -4.055 | 255.2 (100%) | 452.3 (2%) | 140.0 (1%) | 196.0 (1%) | 690.5 (5%) |
| PG(30:0) | 693.4707 | 3.893 | 241.2 (100%) | 153.0 (3%) | 377.2 (1%) | 469.3 (1%) | 693.5 (16%) |
| PE(33:1) | 702.5074 | -2.562 | 241.2 (62%) | 253.2 (9%) | 255.2 (13%) | 281.2 (24%) | 702.5 (100%) |
| PE(33:0) | 704.5231 | -6.387 | 241.2 (7%) | 255.2 (76%) | 269.2 (38%) | 466.3 (5%) | 704.5 (100%) |
| PE(34:1) | 716.5231 | 0.000 | 255.2 (100%) | 281.2 (40%) | 478.3 (2%) | 140.0 (1%) | 716.5 (9%) |
| PG(32:1) | 719.4868 | 12.509 | 227.2 (100%) | 281.2 | 253.2 | 255.2 | 719.5 |
| PG(32:0) | 721.502 | -3.188 | 255.2 (100%) | 153.0 (4%) | 391.2 (2%) | 483.3 (2%) | 721.5 (23%) |
| PE(35:1) | 730.5392 | -3.696 | 269.2 (100%) | 281.2 (41%) | 255.2 (9%) | 478.3 (3%) | 730.5 (15%) |
| PG(33:1) | 733.502 | -0.818 | 241.2 (100%) | 281.2 (50%) | 255.2 (17%) | 253.2 (12%) | 733.5 (40%) |
| PG(33:0) | 735.5177 | -6.662 | 241.2 (3%) | 255.2 (33%) | 269.2 (16%) | 405.2 (2%) | 735.5 (100%) |
| PG(34:2) | 745.5025 | -1.610 | 253.2 (100%) | 281.2 (36%) | 153.0 (5%) | 417.2 (2%) | 745.5 (32%) |
| PG(35:2) | 759.5181 | -1.975 | 253.2 (11%) | 267.2 (25%) | 281.2 (12%) | 295.3 (5%) | 759.5 (100%) |
| PG(35:1) | 761.5333 | -2.495 | 269.2 (100%) | 281.2 (51%) | 255.2 (25%) | 241.2 (16%) | 761.5 (46%) |
| PG(35:0) | 763.5283 | 4.060 | 269.2 (31%) | 405.2 (20%) | 676.5 (44%) | 762.5 (16%) | 763.5 (100%) |

**Table S4:**
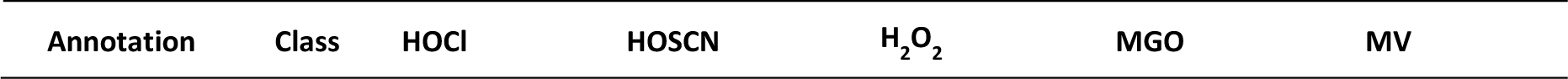
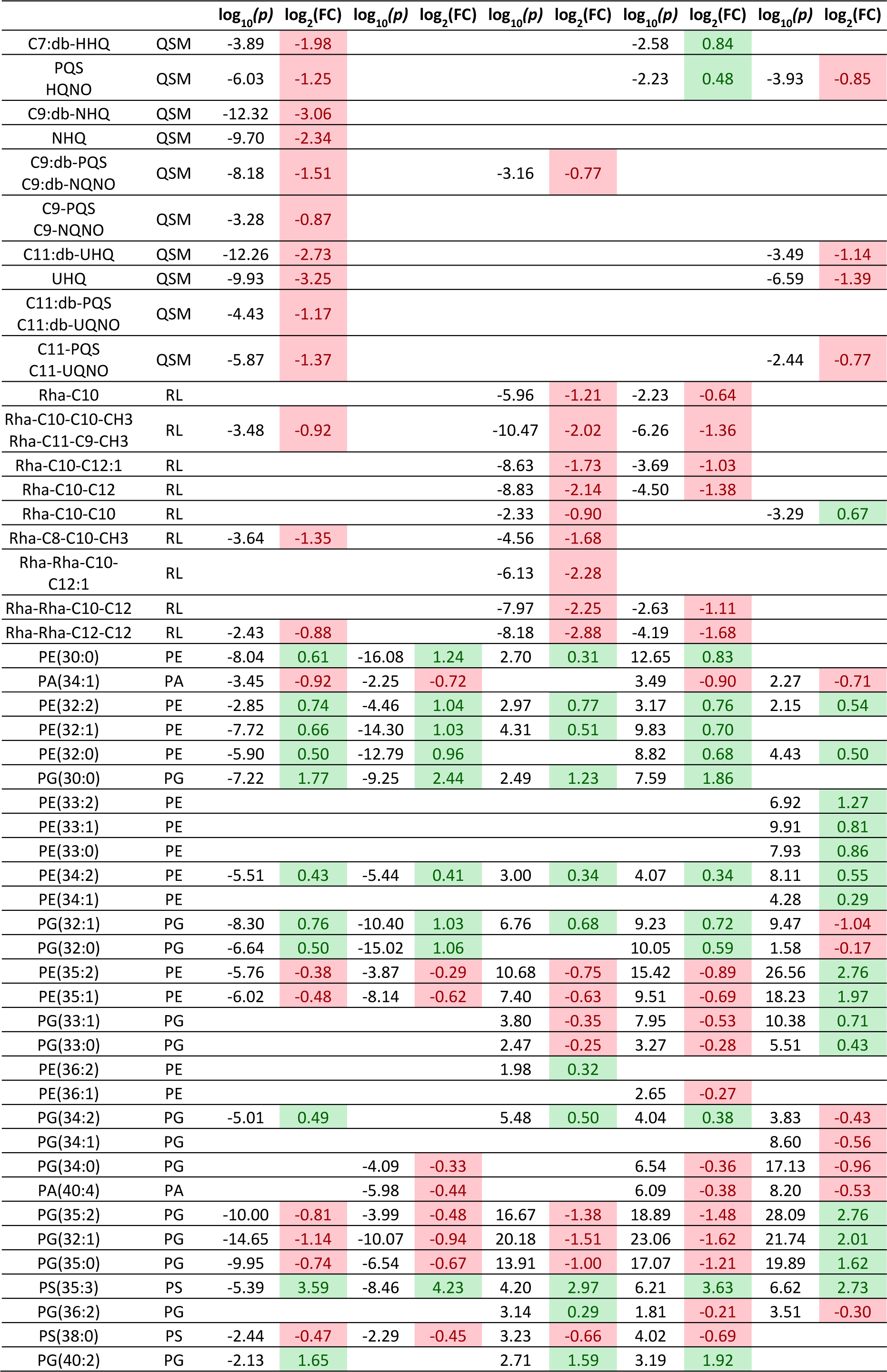
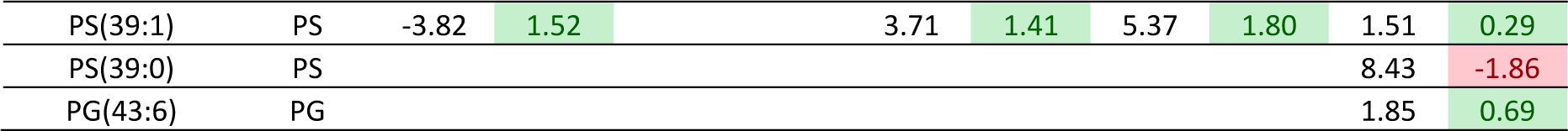
Fold change in intensity and the associated p value for each identified metabolite when exposed to each oxidant. P values were calculated using Students t-test, with a threshold of 0.05. Fold change is calculated using the absolute values of change between the two group means, prior to normalisation.

**Table S5:**
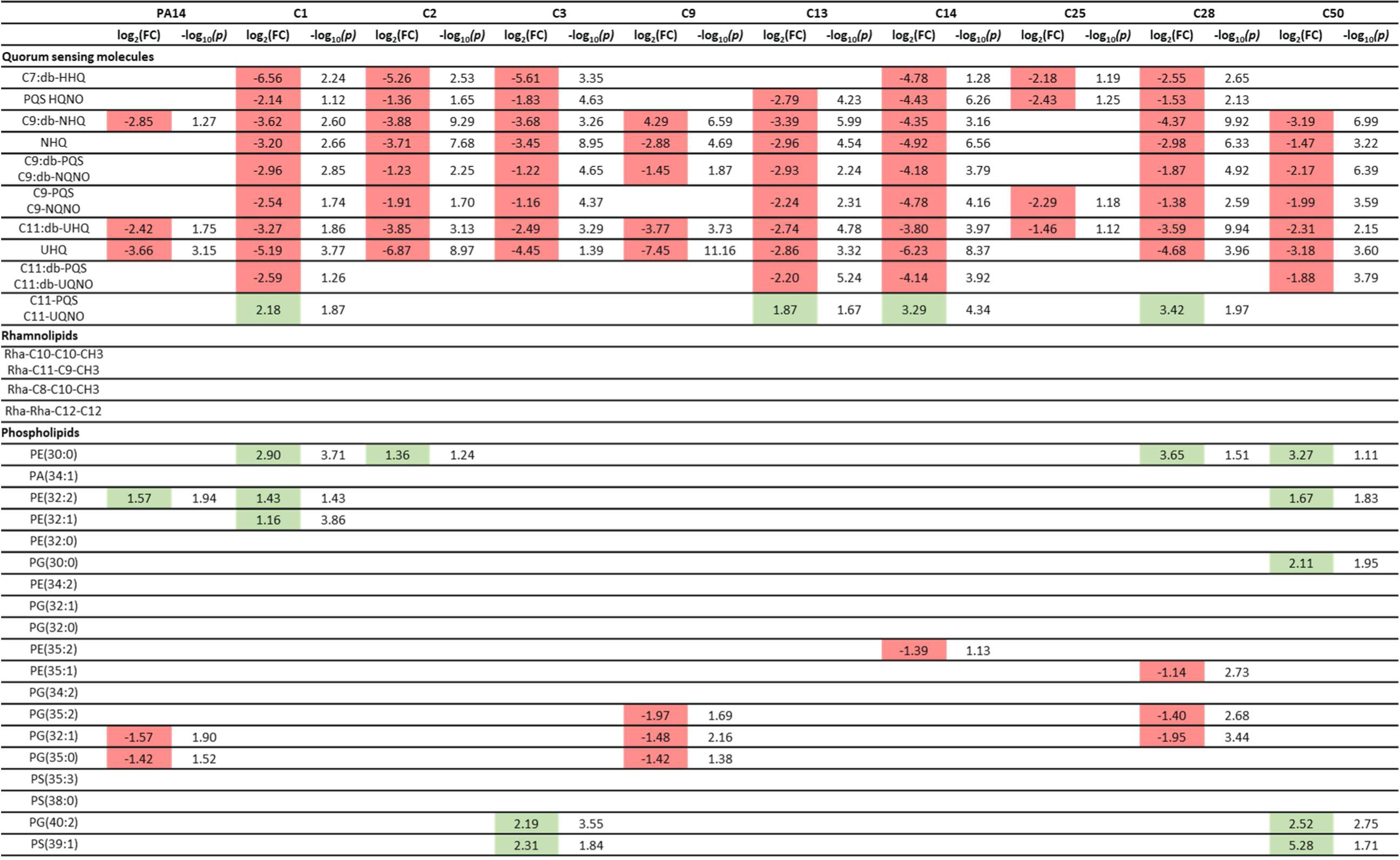
Fold change in intensity and the associated p value for each of the HOCl biomarkers established in PA14 in clinical isolates grown in HOCl. P values were calculated using Students t-test, with a threshold of 0.05. Fold change was calculated using the absolute values of change between the two group means, prior to normalisation.

**Table S6:**
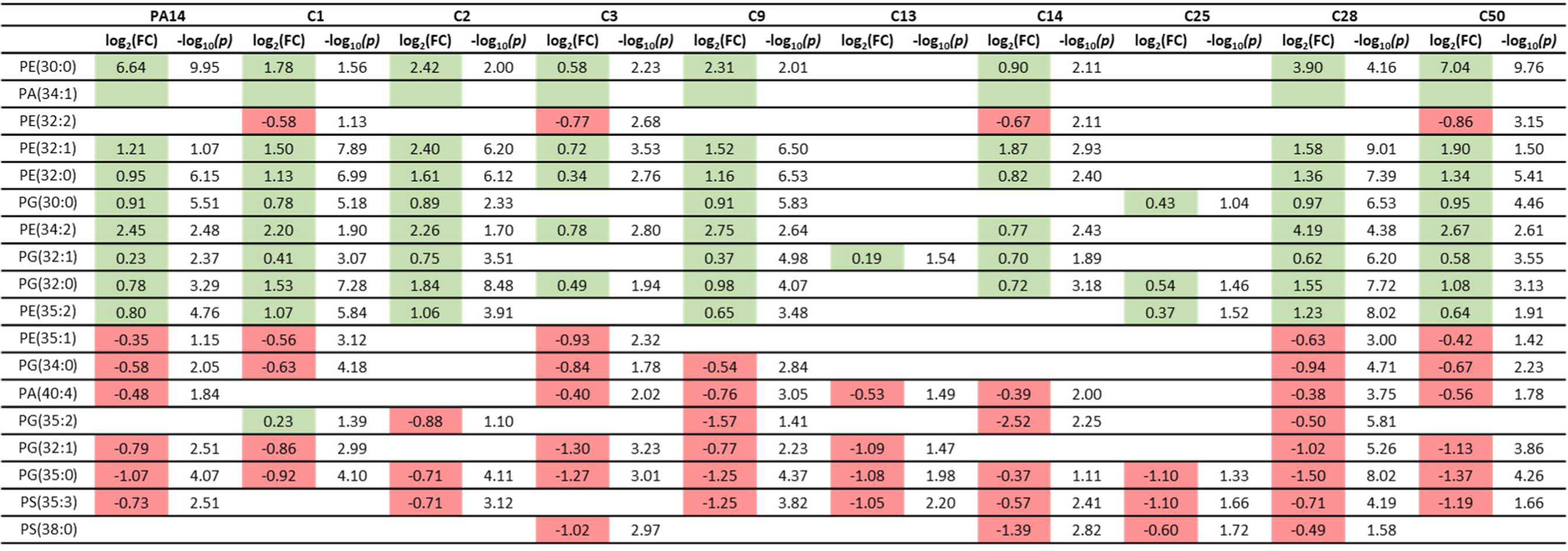
Fold change in intensity and the associated p value for each of the HOSCN biomarkers established in PA14 in clinical isolates grown in HOSCN. P values were calculated using Students t-test, with a threshold of 0.05. Fold change was calculated using the absolute values of change between the two group means, prior to normalisation.

